# Stress-Responsive Protein IFRD1 Protects Assembled Ribosomes via a Ribosome-Salvaging Mechanism

**DOI:** 10.64898/2026.05.03.720925

**Authors:** Charles J. Cho, Molly K. Crowder, Amala K. Rougeau, Thanh Nguyen, Steven J. Bark, Sammy Lee, Jeffrey W. Brown, Jason C. Mills

**Affiliations:** Division of Digestive Diseases, Department of Medicine, Emory University, Atlanta, GA; Graduate Program in Genetics and Molecular Biology, Emory University, Atlanta, GA; Graduate Program in Biochemistry, Cell and Developmental Biology, Emory University, Atlanta, GA; Section of Gastroenterology and Hepatology, Department of Medicine, Baylor College of Medicine, Houston, TX; Cancer & Cell Biology program, Graduate School of Biomedical Sciences, Baylor College of Medicine, Houston, TX; Division of Gastroenterology, Department of Medicine, Washington University in St. Louis, School of Medicine, St. Louis, MO; Program of Molecular and Cell Biology, Washington University in St. Louis, School of Medicine, St. Louis, MO; Program of Biochemistry, Biophysics, and Structural Biology, Washington University in St. Louis, School of Medicine, St. Louis, MO; Program of Developmental, Regenerative, and Stem Cell Biology, Washington University in St. Louis, School of Medicine, St. Louis, MO; Department of Pathology & Immunology, Baylor College of Medicine, Houston, TX; Department of Molecular and Cellular Biology, Baylor College of Medicine, Houston, TX

**Keywords:** IFRD1, ribosome, injury, salvaging, paligenosis

## Abstract

The ability of epithelial cells to cope with injury and undergo regeneration depends on tightly coordinated cellular responses. IFRD1 is a stress-responsive protein that is evolutionarily conserved and required for the cellular regeneration program paligenosis; however, how IFRD1 works in paligenosis is not known. Here we demonstrate that IFRD1 is primarily a cytosolic ribosome-binding protein, specifically binding 80S monosomes that are not actively engaged in translation. Using multiple in vivo and in vitro injury models, including cerulein-induced pancreatitis in mice and tunicamycin-induced ER stress in cell culture, we demonstrate that IFRD1 acts as a ribosome-salvaging factor, preventing ribosomes from degradation. In the absence of IFRD1 during ER stress, non-translating 80S ribosomes were unstable and prone to disassembly and selective degradation. The resulting accumulation of degraded ribosomal subunits overwhelmed cellular autophagic machinery, as evidenced by accumulation of the autophagy-tagging protein p62, even though overall autophagic flux remained unaffected. Ultimately, cells lacking IFRD1 showed reduced mTORC1 activity followed by increased cell death, consistent with patterns observed in cells lacking IFRD1 during paligenosis. Thus, we detail a previously unrecognized cellular function for IFRD1 in stabilizing and preserving the mature ribosome pool during metabolic and translational transitions such as paligenosis.

## Introduction

After injury, epithelial cells undergo dynamic changes to cope with cellular damage and facilitate regeneration. These recovery processes are delicately coordinated and require both cell-intrinsic and -extrinsic responses. Key signaling shifts during this transition include the activation of immediate early genes that govern stress-induced gene expression, the disruption of energy-sensing pathways (e.g., AMPK and mTOR), and the phosphorylation of eIF2α to slow down global protein translation^1-4^. The effectors of the eIF2α phosphorylation, which blocks ribosomal protein translation, include the Activating Transcription Factor (ATF) family of transcriptional regulators (e.g., ATF3 and ATF4) and DDIT3 (CHOP)^5-7^, while other such effectors are still being elucidated.

Recently, interferon-related developmental factor 1 (IFRD1) was identified as a key injury-responsive protein whose mRNA and protein expression increases when eIF2α is phosphorylated^8-11^. IFRD1 is remarkably well conserved throughout biology, with stress-induced functions identified across eukaryotic kingdoms from yeast to plants to insects to mammals^12-15^. Multiple physiological roles for IFRD1 have been identified across various cellular contexts; and in all but a few cases, the role for IFRD1 is in response to stress, injury and/or in regeneration. For example, IFRD1 influences the severity of lung inflammatory disease in patients with cystic fibrosis through modulating neutrophil function and skeletal muscle regeneration by regulating MyoD and NF-κB^16,17^. Furthermore, IFRD1 is essential for secretory cells in the mammalian stomach corpus and pancreas to respond to tissue damage via paligenosis^12^, and it is required for paligenosis-like response of *Drosophila* enterocytes to tissue and stem cell injury in the gut and during recruitment of intestinal stem cells after surgical resection^14,18^. IFRD1 also has a role in response to viral infection^19^. One notable exception to this rule is observed in the bladder where IFRD1 is expressed under steady-state conditions and contributes to epithelial homeostasis, as evidenced by aberrant bladder epithelium characteristics in *Ifrd1-/-*mice^10^.

Although the physiological role of IFRD1 in stress response and regeneration has consistently been observed, the molecular function of this protein remains unclear. Several studies have implicated IFRD1 as a direct or indirect modulator of histone deacetylases (HDACs), working to regulate NF-kB transcriptional activity by modulating its acetylation and potentially working to affect transcription of diverse other genes by epigenetic modification of histones^17,19-22^.

Here, we report that IFRD1 is a cytosol-enriched protein that binds specifically to 80S ribosomes that are not part of polysomes (i.e., multiple ribosomes translating the same mRNA), thus preventing ribosomes from degradation during cellular stress. These findings establish IFRD1 as a key factor in ribosome-salvaging process, a newly defined cellular function that appears to be critical across multiple eukaryotic kingdoms.

## RESULTS

### IFRD1 is a general injury-responsive protein in vivo and in vitro

To understand the mechanisms governing IFRD1 function, we examined multiple cellular/tissue injury models, both in vivo and in vitro. In particular, we focused on a model of acute pancreatitis, in which cerulein injury induces pancreatic acinar cells to undergo paligenosis^12^. As previously described, *Ifrd1* mRNA and protein increase when cells experience decreased global translation, allowing IFRD1 to be active in an injury context where, for example, ER is under stress^8-11^. Specifically, global suppression of translation via eIF2α phosphorylation at Ser51 paradoxically stabilizes *Ifrd1* mRNA by promoting IFRD1 protein translation via its major open reading frame (ORF) rather than an abortive upstream ORF^8,9,11^. Within one hour of a single cerulein injection intraperitoneally in mice, we observed an abrupt, near-complete reduction in global translation (Figure 1A), as measured by incorporation of the aminoacyl-tRNA analogue puromycin. Acinar cells are more translationally active than the other principal pancreatic cell types (i.e., islet or ductal cells) under baseline conditions (Figure 1B), and the translational halt was most prominent in acinar cells (Figure 1C), which are the cell type that undergoes paligenosis in the cerulein injury model. This translational halt coincided with significantly increased eIF2α phosphorylation (p-eIF2α) within 30 minutes (Figure 1D). As expected, IFRD1 protein levels did not immediately change, but did eventually accumulate robustly by one hour after injury (Figure 1E, left and middle), even as p-eIF2α returned to baseline (Figure 1F). *Ifrd1* mRNA also increased significantly by 1 hour (Figure 1E, right), consistent with the previously demonstrated mechanism^9^.

**Figure 1.**
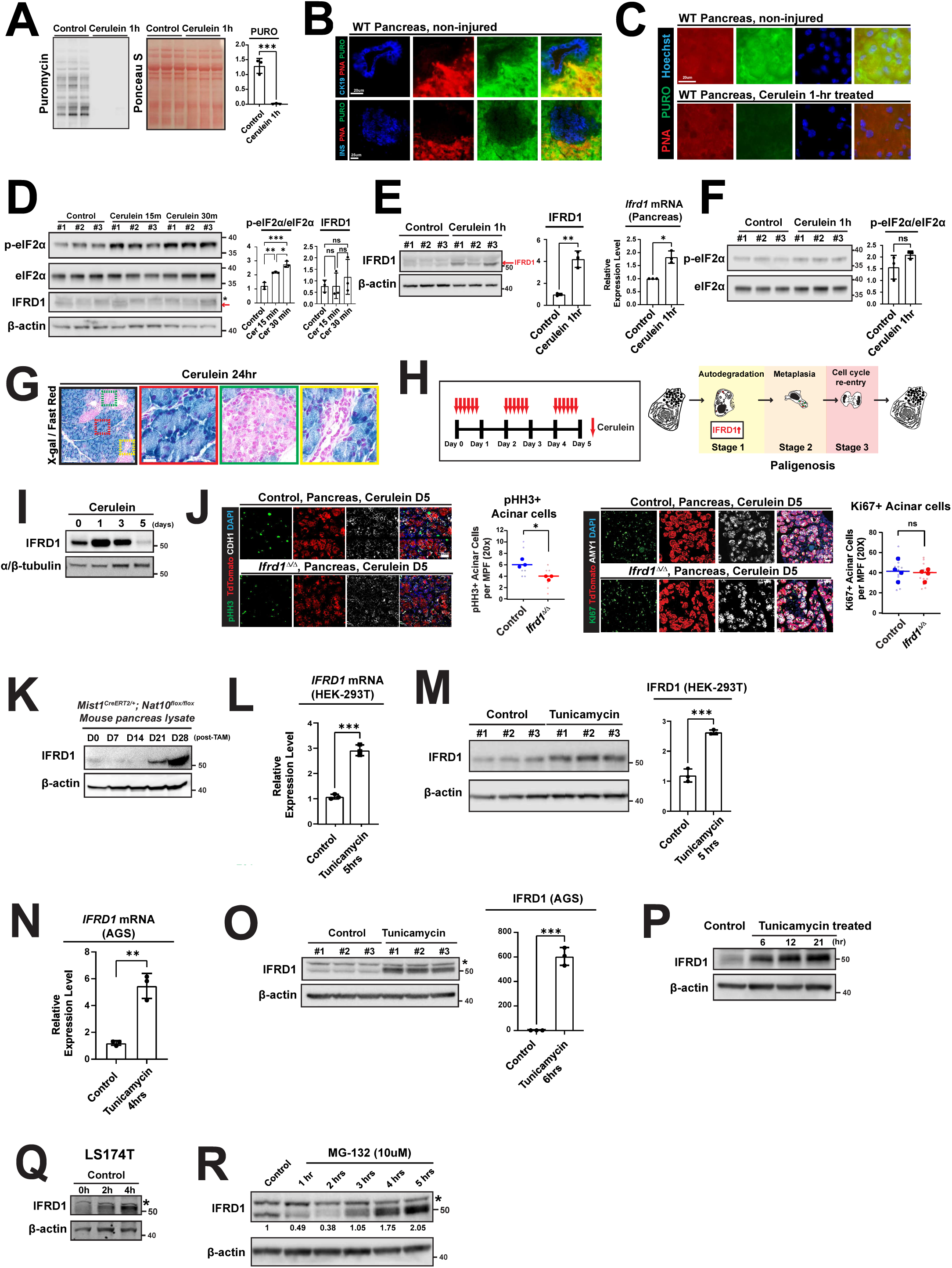

IFRD1 expression was exclusive to acinar cells during injury. Because IFRD1-specific, sensitive antibodies have still not been developed to stain endogenous IFRD1 in mouse tissue sections^10^, we took advantage of the lacZ allele in the *Ifrd1* null allele to do an enzymatic stain for expression from the *Ifrd1* promoter (Figure 1G). Using a prolonged cerulein protocol that we previously have shown causes acinar cells to undergo all the three stages of paligenosis (Figure 1H), we observed a steady increase in IFRD1 peaking at 1 day (Figure 1I). Consistent with previous global knockout studies, specifically deleting *Ifrd1* in acinar cells using a conditional knockout model (*Ifrd1^flox/flox^; Rosa26^LSL-TdTomato;^ Mist1^CreERT2/+^*), when compared to control (*Ifrd1^flox/+^; Rosa26^LSL-TdTomato;^ Mist1^CreERT2/+^*) resulted in a significant reduction in cells progressing to stage 3 of paligenosis (the stage where paligenotic cells undergo mitosis). Specifically, we noted a decrease in TdTomato-positive cells re-entering M-phase (determined by pHH3 positivity; 6.2 ±0.7 vs. 3.8 ±0.4 /MPF, *P* = 0.012, Figure 1J, left), despite a similar number of cells having at least entered G1 of the cell cycle (Ki-67 positive; 41.9 ±9.6 vs. 38.6 ±5.5 /MPF, *P* = 0.57, Figure 1J, right). The discrepancy between M-phase and total cell cycle effects from loss of *Ifrd1* are consistent with our previous work showing that the checkpoint for stage 3 of paligenosis involves mTORC1 activation to drive cells into S-phase. Loss of IFRD1—just like the mTORC1 inhibitor rapamycin—blocks the stage 3 mTORC1 activation, causing cells to arrest in the G1 phase of the cell cycle, where they express Ki-67 but not S- or M-phase markers^12^.

We next looked at mice with acinar cells lacking NAT10, which is required for 18S rRNA acetylation and ribosome biogenesis, because we previously showed NAT10 and ribosome biogenesis were also required for maintaining acinar cell identity and preventing apoptotic cell death^23^. Over time, pancreata with acinar cells lacking *Nat10* accumulated IFRD1, suggesting IFRD1 might be sensitive to changes in ribosome census (Figure 1K), an observation we will return to in experiments to be described below.

Next, we wanted to establish in vitro systems where we could study the mechanisms of IFRD1’s role in the injury response. We leveraged the tunicamycin injury model, which is widely used in the field to induce ER stress with consequent triggering of eIF2α phosphorylation and translational inhibition; IFRD1 has also been previously shown to accumulate after tunicamycin^9,12^. In HEK-293T cells, we observed a sharp increase in IFRD1 mRNA and protein within 5 hours (Figures 1L–M). Similar results were obtained in AGS (gastric adenocarcinoma) and LS174T (colon cancer) cell lines (Figures 1N–Q). Finally, treating AGS cells with the proteasome inhibitor MG-132—which also triggers eIF2α phosphorylation in long-term treatment^24^—resulted in increased IFRD1 levels (Figure 1R). Collectively, these data demonstrate that IFRD1 is a conserved stress-responsive protein across various cell types and injury contexts, particularly in settings of global protein translation block. Furthermore, these results provide a robust platform for investigating its mechanistic function.

### IFRD1 is an 80S monosome-binding protein

To elucidate the biological function of IFRD1, we used an unbiased immunoprecipitation (IP) approach followed by mass spectrometry to determine the IFRD1 interactome (i.e., the proteins that IFRD1 binds). Initially, we pulled down IFRD1 using an anti-IFRD1 antibody in LS-174T cells after treatment with tunicamycin for 3.5 and 20 hours. PANTHER analysis revealed a high correlation in Gene Ontology gene sets between these two timepoints; both timepoints showed significant enrichment in pathways related to ribosomes and translation (Figure 2A, left and center, see Supplementary Table 2 for the complete list).

**Figure 2.**
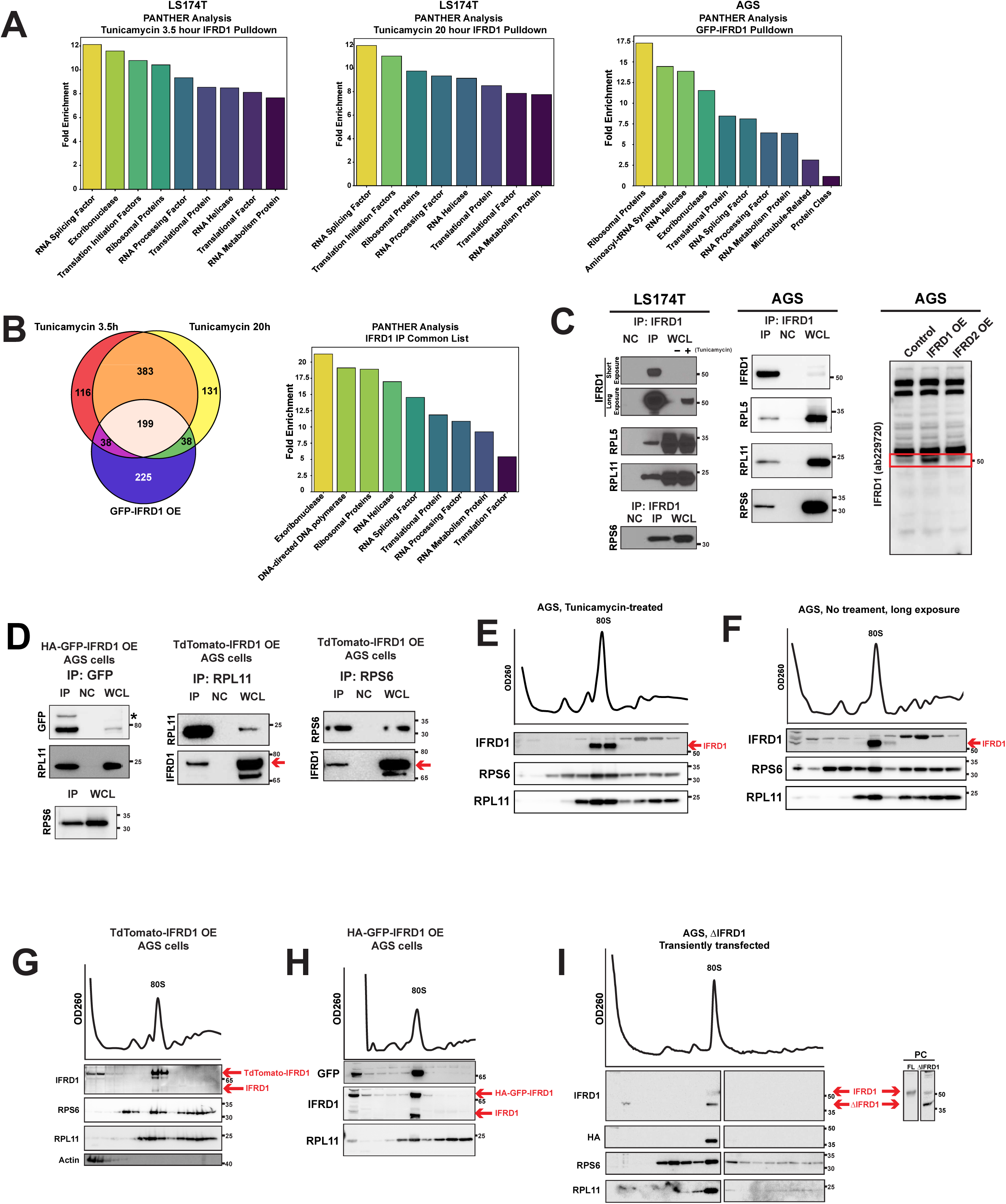

To control for potential cross-reactive binding of the anti-IFRD1 antibody to other proteins, we conducted a complementary experiment with anti-GFP immunoprecipitation (IP) from AGS cells stably overexpressing HA-GFP-tagged IFRD1. Using this approach, the captured proteins were still enriched in ribosome-and translation-related pathways (Figure 2A, right). Across all three IFRD1 IP datasets, we identified a relatively large, consistent, unbiased interactome of 199 proteins (Figure 2B, left). Again, as might be predicted, PANTHER analysis of this core interactome revealed that eight of the top nine enriched candidates were ribosomal components (Figure 2B, right). In all experimental sets, we observed robust enrichment of IFRD1 in the pulldown list, while the paralog IFRD2 was consistently undetected, confirming the specificity of our immunoprecipitation and the unique interactome of IFRD1.

Our results were initially surprising because IFRD1 has previously been described as a nuclear protein that interacts with chromatin^17,21,22^, and we did not observe the enrichment of chromatin remodeling proteins, including HDACs, KATs, and sirtuins; however, weak pulldown of SIN3A and HDAC1 was detected in two out of three independent experimental sets (Supplementary Table 2). Ribosomes are abundant and can thus be prone to nonspecific pulldown^25^. So, to further validate our findings, we performed IPs in AGS cells, our chosen in vitro injury model, due to their rapid and dramatic response to tunicamycin (Figures 1N & 1O). Using the same rabbit polyclonal antibody as the initial discovery set, we observed a robust interaction between IFRD1 and the large ribosomal subunit components RPL5 and RPL11, as well as the small subunit component RPS6, following tunicamycin injury in both the LS174T cell line (Figure 2C, left) and AGS cell line (Figure 2C, middle). We have also validated that the antibody does not recognize IFRD2, a paralog of IFRD1 with a similar size and high sequence similarity that could confound our data (Figure 2C, right).

The interaction between IFRD1 and ribosomal proteins was further confirmed using an anti-GFP antibody in AGS cells overexpressing C-terminus tagged HA-GFP-IFRD1 (Figure 2D, left), demonstrating that the binding was not an artifact of the anti-IFRD1 antibody. Additionally, a reciprocal pulldown using anti-RPL11 and anti-RPS6 antibodies confirmed the IFRD1-ribosomal protein interaction in cells transfected with N-terminus TdTomato-tagged IFRD1 (Figure 2D, middle & right).

While these data provide evidence that IFRD1 is a ribosome-binding protein, they do not specify whether it interacts directly with isolated, individual ribosomal proteins, with large or small ribosomal subunits, or with assembled 80S ribosome structures such as monosomes or polysomes. Thus, we next used sucrose density gradient centrifugation to determine the specific ribosomal assembly to which IFRD1 binds. After tunicamycin injury, we found IFRD1 was associated specifically with monosomes and not with other ribosomal components, such as polysomes or individual large and small subunits (Figure 2E). This demonstrates that IFRD1 binds intact ribosomes, not just individual ribosomal proteins and explains why the IFRD1 interactome is large, given how many proteins are associated with ribosomes^26^. We also confirmed that IFRD1 was exclusively bound to monosomes in non-injured cell lysates (Figure 2F), as well as in cells overexpressing N-terminally TdTomato-tagged IFRD1 (Figure 2G) or C-terminally HA-GFP-tagged IFRD1 (Figure 2H) without tunicamycin injury. These results suggest that the monosome-binding property of IFRD1 is an inherent characteristic of the protein rather than a state acquired following injury. Although IFRD1 lacks canonical or previously-described RNA-binding motifs, it contains an intrinsically disordered region (IDR) at its N-terminus. Because IDRs can facilitate versatile binding to mRNAs or other structures^27^, we investigated whether this region is necessary for ribosome interaction by generating a truncated version of IFRD1 lacking the IDR (ΔIFRD1, lacking amino acids 1–69). Interestingly, the deletion of the IDR did not abolish the binding of ΔIFRD1 to monosomes, indicating that this domain is dispensable for the IFRD1-ribosome interaction (Figure 2I). In summary, IFRD1 is a ribosome-binding protein that specifically interacts with 80S monosomes.

### IFRD1 localizes to cytosol and is excluded from stress granules

Ribosome-binding proteins can localize to multiple cellular compartments, as the ribosome life cycle spans the nucleolus, nucleoplasm, and cytoplasm. This localization can help infer the function of specific ribosome-binding proteins; for example, presence in the nucleolus suggests a role in ribosome biogenesis, whereas proteins that interact with mature ribosomes are typically enriched in the cytoplasm, where fully assembled 80S ribosomes reside. We therefore investigated the subcellular localization of IFRD1 using immunofluorescence (IF) imaging of TdTomato-tagged IFRD1, which we previously demonstrated functions just as endogenous IFRD1 in the monosome-binding assay (Figure 2F). The TdTomato signal was primarily observed in the cytoplasm (Figure 3A) and was almost entirely absent from the nucleus (i.e., no overlap with DAPI staining). Furthermore, nucleolin (NCL)-positive nucleoli were devoid of IFRD1 signal (Figure 3B). Corroborating the imaging data, subcellular fractionation—with compartment-specific markers including Histone H3 (nucleus), and the p110 isoform of ADAR1 (nucleus, especially the nucleolus)—showed that IFRD1 was predominantly in the cytoplasmic fraction (Figure 3C). Finally, the cytoplasmic IFRD1 signal overlapped highly with all four ribosomal proteins examined: RPS6, RPL13, RPL5, and RPL11 (Figure 3D). These data are consistent with a model in which IFRD1 primarily binds to mature, assembled 80S ribosomes, which are localized primarily in the cytoplasm.

**Figure 3.**
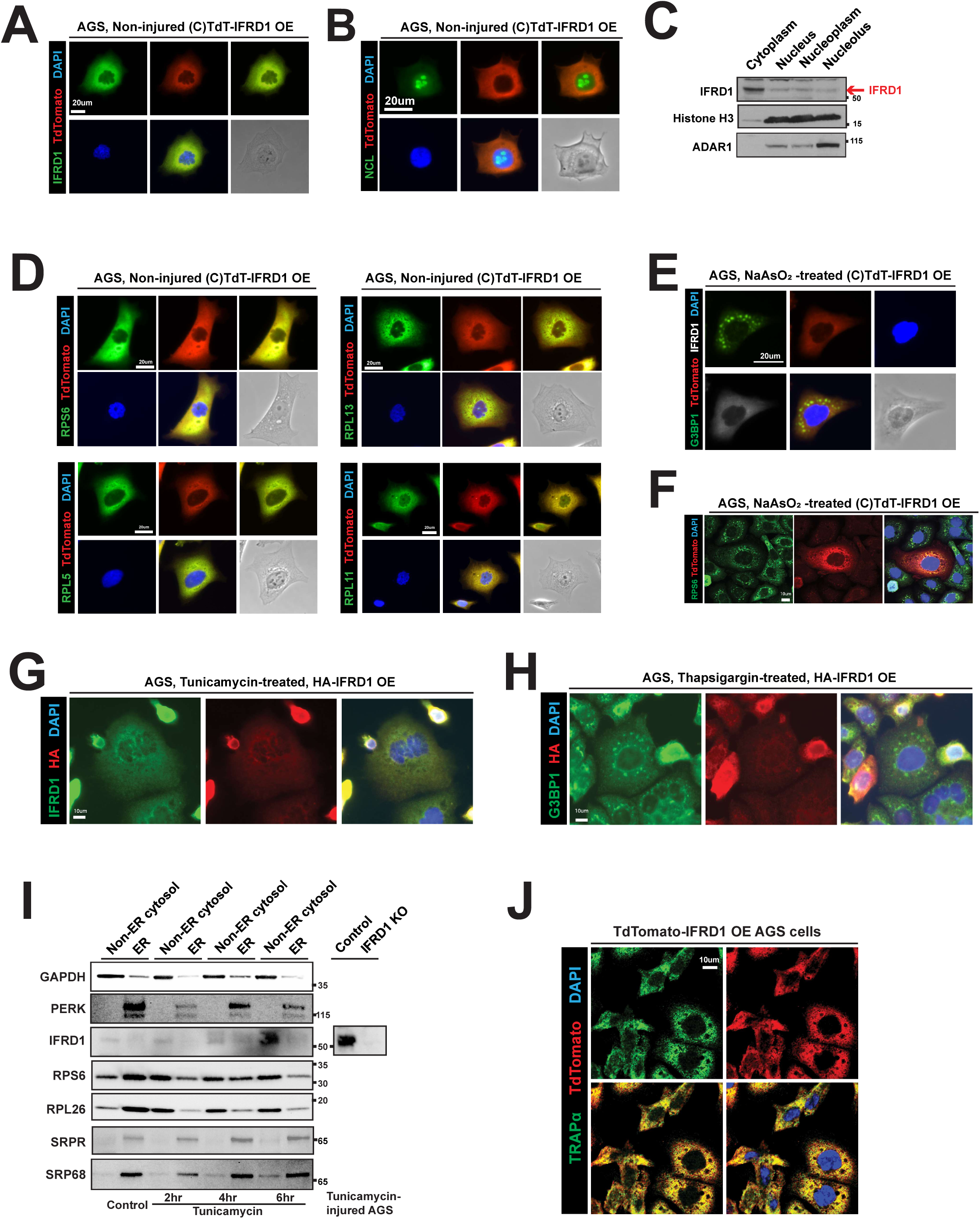

We next investigated if IFRD1 localized to stress granules (SGs), as these membrane-less organelles are, like IFRD1, also induced upon eIF2α phosphorylation and global translation block. We treated cells with sodium arsenite (NaAsO_2_) to induce SGs within 30 minutes^28^. Interestingly, the TdTomato-IFRD1 signal was not enriched in G3BP1-positive SGs (Figure 3E), remaining in the cytoplasm. When RPS6 puncta were used as a marker for SGs, there was some colocalization with IFRD1, but the vast majority of IFRD1 was not in SGs (Figure 3F). Transfection of HA-tagged IFRD1 in cells also showed a generalized, reticular cytoplasmic distribution, overlapping with an antibody-recognizing endogenous IFRD1 (Figure 3G). When HA-IFRD1-transfected cells were induced to form SGs via thapsigargin, HA-tagged IFRD1 also did not accumulate in G3BP1 puncta/SGs (Figure 3H). SGs are typically composed of pre-initiation complexes (i.e., 40S subunits bound to mRNA and translation initiation factors) and specific marker proteins like G3BP1; they usually exclude the 60S subunit. Thus, the lack of IFRD1 localization to SGs again supports the interpretation that IFRD1 binds assembled ribosomes that remain in the cytosol rather than ribosomal 40S subunits sequestered in a halted translation state within SGs. Finally, because monosomes can exist in either free-floating cytosolic or ER-bound forms, we performed digitonin-based fractionation and a ±tunicamycin injury time course to determine the subcellular distribution of IFRD1. We found that the majority of IFRD1 can be found in the non-ER cytosolic fraction, although at the 4-hour timepoint, some IFRD1 was ER-associated (Figure 3I).

Immunofluorescence staining revealed that the IFRD1 signal partially overlapped with TRAPα, an ER-marker (Figure 3J). Taken together, these data indicate that IFRD1 primarily interacts with mature, assembled 80S ribosomes in the cytoplasm, although its interaction with ribosomes on the rough ER may also occur.

### IFRD1 attenuates global translation by binding to a pool of idling ribosomes

We next focused on elucidating the functional consequences of IFRD1 binding to the ribosome. Published cryo-electron microscopy (cryo-EM) data for the IFRD1 paralog IFRD2 in rabbit reticulocyte lysates show that these proteins occupy the P and E sites of the ribosome^29^, as well as the mRNA exit tunnel physically plugged by the IFRD2 protein’s C-terminus, which is also observed in the IFRD1 ortholog (dIFRD1) in Drosophila^30^. Because we have shown here that IFRD1 binds monosomes, it would make sense that mammalian IFRD1—which is structurally similar to IFRD2 and dIFRD1 in the region where those latter two proteins localize—would be associated with monosomes that are “naked”. In other words, mRNA and the IFRD1 protein should be mutually exclusive sterically in ribosomes, and IFRD1 likely is bound to vacant, non-translating ribosomes, which could serve to regulate ribosome availability for translation. In this model, the influence of IFRD1 on global translation would be determined by: 1) the total abundance of IFRD1—binding to ribosomes in a 1:1 ratio; and 2) the duration of IFRD1 binding to ribosomes (i.e., on and off rates).

The net effect of IFRD1 might be to sequester a pool or subset of ribosomes, preserving them from disassembly and also limiting their immediate availability for translation. An open question is whether IFRD1 would need to remain bound to these naked monosomes to keep them from disassembly or if, instead, an IFRD1 protein can move from ribosome to ribosome, stabilizing multiple ribosomes. This question of stoichiometry is important because, even after injury, IFRD1 seems to work in diverse cells without being particularly abundant. In any case, from a translational blockade perspective, IFRD1’s function in generating and/or stabilizing a pool of non-translating ribosomes would represent a different mechanism in terms of modulation of global protein translation relative to other mechanisms like phosphorylation of eIF2α or loss of mTORC1 activity with dephosphorylation of 4E-BP1.

We first investigated whether CRISPR-mediated knockout or overexpression of IFRD1 altered global translation levels using puromycylation assays. In the tunicamycin-induced injury model in AGS cells, we observed a substantial reduction in global translation (Figure 4A,^31^), which was also noted following translation elongation inhibitor, cycloheximide (CHX) treatment. On the other hand, we did not observe a significant difference in translation levels between IFRD1-knockout or overexpressed cells vs. control cells under stress (Figure 4B). This aligns with the idea that IFRD1 binds primarily to non-translating ribosomes and therefore has a limited impact on global translation compared to active inhibitors of translation initiation, such as phosphorylation of eIF2α^31^.

**Figure 4.**
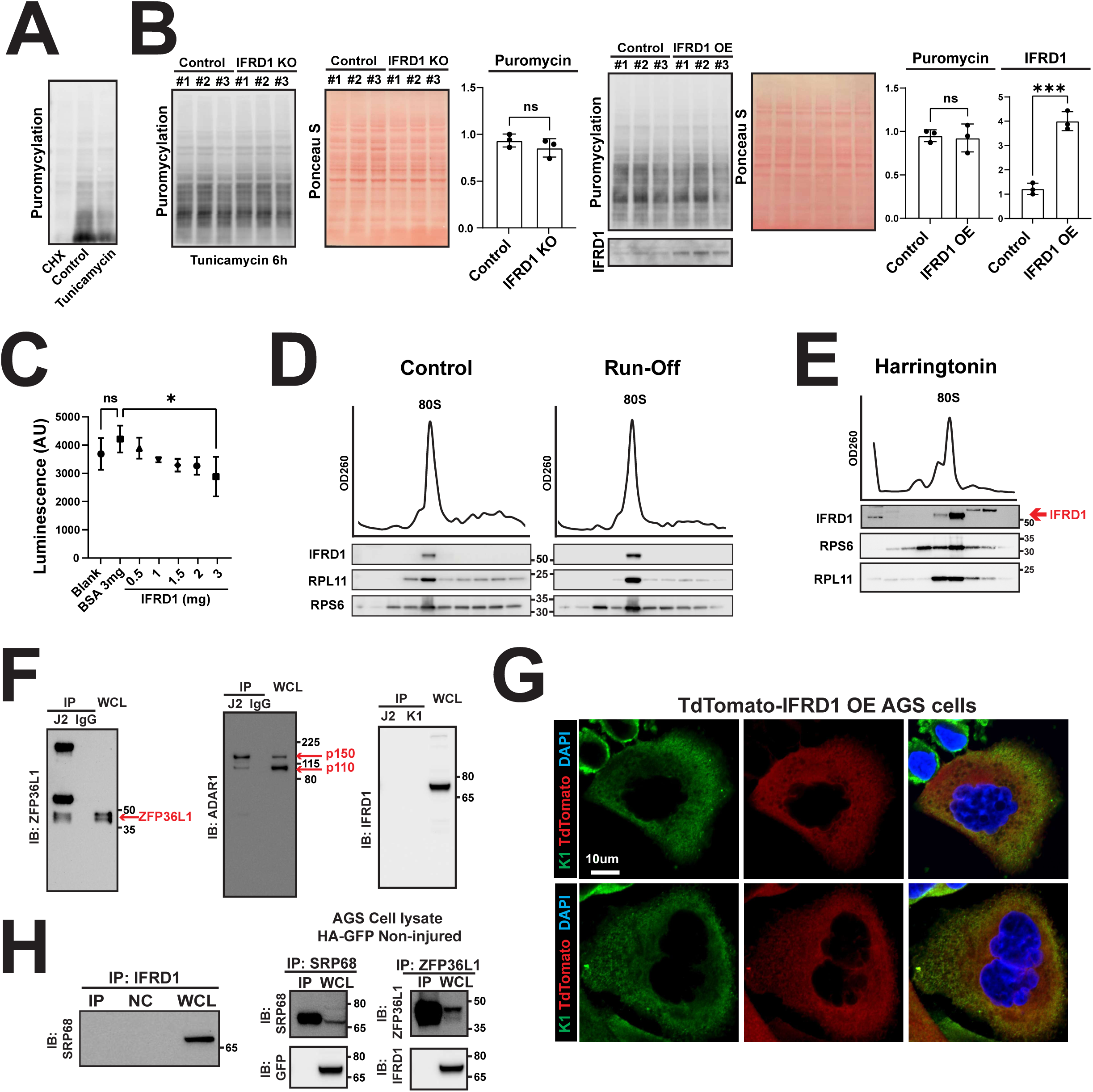

To enhance our sensitivity for detecting subtle changes in translation that might be missed in the generalized assessment of translation across a population of cells using puromycylation, we leveraged a rabbit reticulocyte lysate (RRL) system that would allow us to titrate IFRD1 abundance in a stripped-down translation assay. We varied concentrations of purified human IFRD1, using BSA as a size-matched negative control and luciferase mRNA as a reporter for translation activity in the RRL. We observed a concentration-dependent, linear reduction in the luciferase signal—reaching a 31.6% decrease at the maximum dose (Figure 4C)—which was absent in the BSA controls. Thus, in a system where IFRD1 abundance is the only variable, we can see that IFRD1 can block translation, likely by reducing the pool of ribosomes available for translation through sequestration and preventing the 80s ribosome from disassembling, a prerequisite for loading mRNA.

We next asked if association of IFRD1 with the monosomes is altered under conditions that disrupt polysome integrity or translation initiation; if IFRD1 binds to non-translating ribosomes, its association should be unaffected by these conditions. Both the addition (Figure 4D, left) and omission (aka, “run-off” experiment, Figure 4D, right) of cycloheximide, as well as stalling ribosomes during translation initiation via harringtonine treatment (Figure 4E), consistently showed IFRD1 binding exclusively to the monosome fraction. This further demonstrates that IFRD1 binding is independent of ribosomes actively engaged in translation. We next asked whether IFRD1 interacts with mRNAs, RNA-binding proteins, or translation-associated complexes, as such interactions would suggest association with translating ribosomes. Using J2 and K1 antibodies that recognize dsRNA, Figure 4F shows that J2 could immunoprecipitate known RNA binding proteins like ZFP36L1 (left) and ADAR1 (middle), but neither J2 nor K1 immunoprecipitated IFRD1 (right). Plus, immunofluorescence for K1 did not show overlap with dsTomato-IFRD1 (Figure 4G).

Additionally, IFRD1 did not bind to SRP68 (Figure 4H, left and middle), a component of the signal recognition particle that targets mRNA to the rER, nor did it bind with the RNA-associated ZFP36L1 (Figure 4H, left). In sum, we found no evidence that IFRD1 is associated with actively translating ribosomes.

### IFRD1 is a ribosome-salvaging protein, the absence of which leads to instability of 80S and 60S ribosomes and inhibition of mTOR signaling

We next considered the possibility that IFRD1 influences ribosomal stability. This hypothesis is supported by the fact that IFRD1 abundance increases during injury of the type that causes degradation of organelles^32^. Given that IFRD1 specifically binds monosomes that are largely not mRNA-associated, we hypothesized that IFRD1 could work to stabilize ribosomes during translation blockade. Thus, IFRD1 may be a ribosome “salvage factor” that stabilizes the monosome state and prevents disassembly. After the monosome splits into 60S and 40S subunits, those can be further disassembled, into individual ribosomal proteins, which can then be degraded. The two subunits are not equally susceptible to such disassembly into constituent individual proteins, nor are those proteins equally susceptible to subsequent degradation. Specifically, the 60S subunit generally is more prone to disassembly than the 40S, and certain 60S (large) ribosomal proteins are particularly prone to subsequent degradation^33,34^.

We assessed the effect of IFRD1 loss after cellular stress on relative ribosomal subunit abundance. Sucrose density gradient centrifugation revealed a substantial reduction in the 60S and 80S fractions, whereas the 40S fraction remained relatively stable in cells either lacking IFRD1 through CRISPR knockout (Figure 5A, left) or stably knocked down using short hairpin RNA (Figure 5A, middle). However, prolonged stress ultimately resulted also in a reduction in 40S ribosomes (Figure 5A, right). These results suggest that assembled 80S ribosomes are prone to disassembly in the absence of IFRD1 and that the 60S large subunit is particularly vulnerable to further destruction.

**Figure 5.**
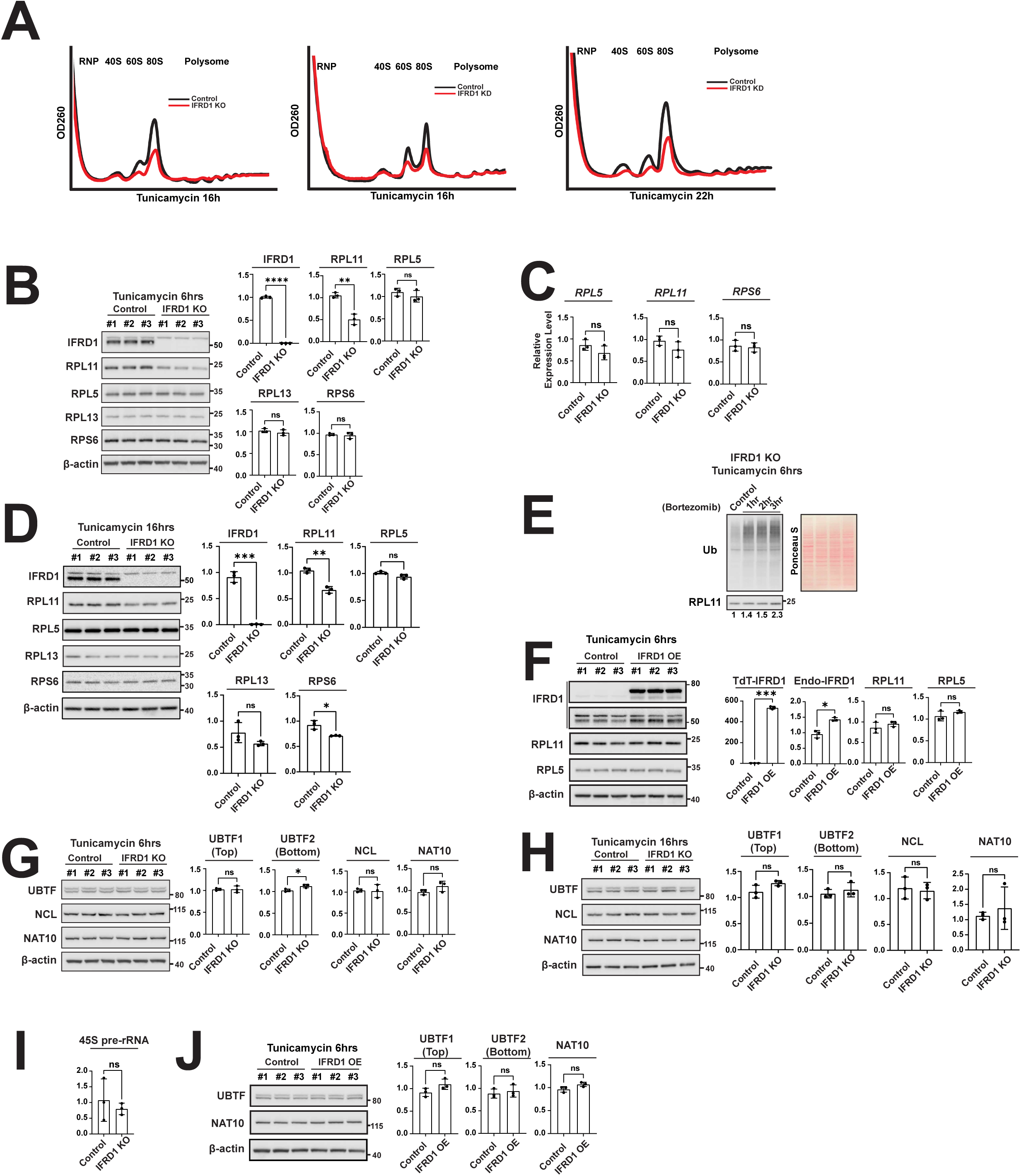

We next examined the abundance of individual ribosomal proteins following injury in the presence or absence of IFRD1. Interestingly, RPL11 levels were significantly decreased within 6 hours of tunicamycin treatment, whereas other large subunit components, such as RPL5 and RPL13, as well as a small subunit component, RPS6, remained unchanged (Figure 5B). The reduction in RPL11 was not driven by transcriptional changes, as we found no significant differences in its mRNA levels or in the transcripts of other ribosomal proteins (Figure 5C). With prolonged tunicamycin treatment, we observed a slight but significant reduction in the small ribosomal subunit RPS6 (Figure 5D), consistent with the reduction in the 60S and 80S peaks (Figure 5A). Nevertheless, the most pronounced decrease was observed for RPL11 (Figure 5D). The selective degradation of specific ribosomal proteins suggests that once monosomes disassemble in the absence of IFRD1, their constituent individual ribosomal proteins may be subject to distinct degradation kinetics or pathways.

Finally, we investigated whether the reduced expression of RPL11 is consistent with UPS-mediated degradation, a well-established mechanism for degrading unassembled ribosomal proteins^33,35^. Indeed, the reduction in RPL11 was ameliorated by bortezomib treatment in tunicamycin-treated IFRD1 knockout samples in a time-dependent manner (Figure 5E).

On the other hand, overexpressing IFRD1 did not result in significant differences in the levels of RPL11 and RPL5 (Figure 5F). The absence of an effect occurred despite a robust increase in TdTomato-tagged IFRD1, the functionality of which we had already validated by demonstrating its ability to bind monosomes (Figure 2G). Thus, IFRD1 does not appear to have some direct effect on ribosomal mRNA transcription or translation, concordant with its preserving ribosomal protein stability in a situation where the proteins are prone to degradation (e.g., during stress-induced ribosomal disassembly).

We next probed the mechanism whereby loss of IFRD1 reduces cellular ability to enter mitosis in response to injury (Figure 1J,^12^). Loss of IFRD1 reduced the monosome pool. Most of these are typically not translating in cells, so we had not previously noticed a major shift in translation activity in cells lacking IFRD1 (Figure 4B). On the other hand, the dramatic loss of the ribosome pool would likely cause a compensatory ribosome biogenesis (RiBi)^36^. However, other than a slight increase in the expression level of UBTF (Upstream Binding Transcription Factor) 2, which is known to have a relatively minor effect on Pol I transcription^37^, all other RiBi-related factors across the process—including UBTF1 (a key RNA polymerase I transcription factor), and NCL and NAT10 (involved in rRNA processing)—remained unaltered 6 hours (Figure 5G) and 16 hours (Figure 5H) after tunicamycin treatment.

Furthermore, 45S pre-rRNA levels remained stable (Figure 5I). Overexpression of IFRD1 also did not alter RiBi (Figure 5J). In sum, a compensatory RiBi response does not occur in the absence of IFRD1 and is therefore unlikely to account for the increased cell death observed in IFRD1 knockout cells.

### A reduction in mTORC1 activity and unregulated autophagy occur in the absence of IFRD1

Autophagy is a conserved stress response activated during various forms of cellular injury. This includes during stage 1 of paligenosis, as seen with the in vivo models (acute pancreatitis and high-dose tamoxifen-induced gastric injury) and in vitro model (tunicamycin) utilized in this study. We investigated whether the loss of IFRD1 altered autophagic flux, potentially rendering cells less fit for survival. In wild-type cells, 6 hours of tunicamycin treatment induced an increase in autophagic flux, characterized by a reduction in p62 levels and the accumulation of the lipidated form of LC3B (LC3B-II), consistent with previous reports (Figure 6A). In contrast, at the same 6-hour time point, IFRD1-deficient cells exhibited a significant accumulation of the autophagy cargo protein p62 compared to wild-type controls. Interestingly, we did not observe a corresponding increase in the accumulation of LC3B-II, which in contrast, trended towards decreased levels (Figure 6B). This pattern of p62 buildup without the expected increase in LC3B-II persisted through 16 hours of prolonged tunicamycin treatment (Figure 6C). To determine if these observations indicated a primary defect in autophagic flux, we challenged the cells with hydroxychloroquine, a pharmacological agent that blocks autolysosomal degradation. Hydroxychloroquine rescued the phenotype of decreased LC3B-II seen in IFRD1-deficient cells with both wildtype and null cells showing accumulation of LC3B-II. Thus, the fundamental autophagic machinery remains functional in the absence of IFRD1. Furthermore, the increase in p62 was not driven by transcriptional changes, as SQSTM1 (p62) mRNA abundance remained unchanged (Figure 6E). Taken together, the data indicate that the observed p62 accumulation was not due to a block in the autophagy pathway itself but potentially due to increase in p62-tagged autophagic substrates. Given how abundant ribosomal proteins are as a fraction of total protein, the observations of significant decreases in monosomes in IFRD1-deficient cells would be expected to increase the burden of degraded ribosomal requiring clearance, potentially placing increased demand on the autophagic pathway.

**Figure 6.**
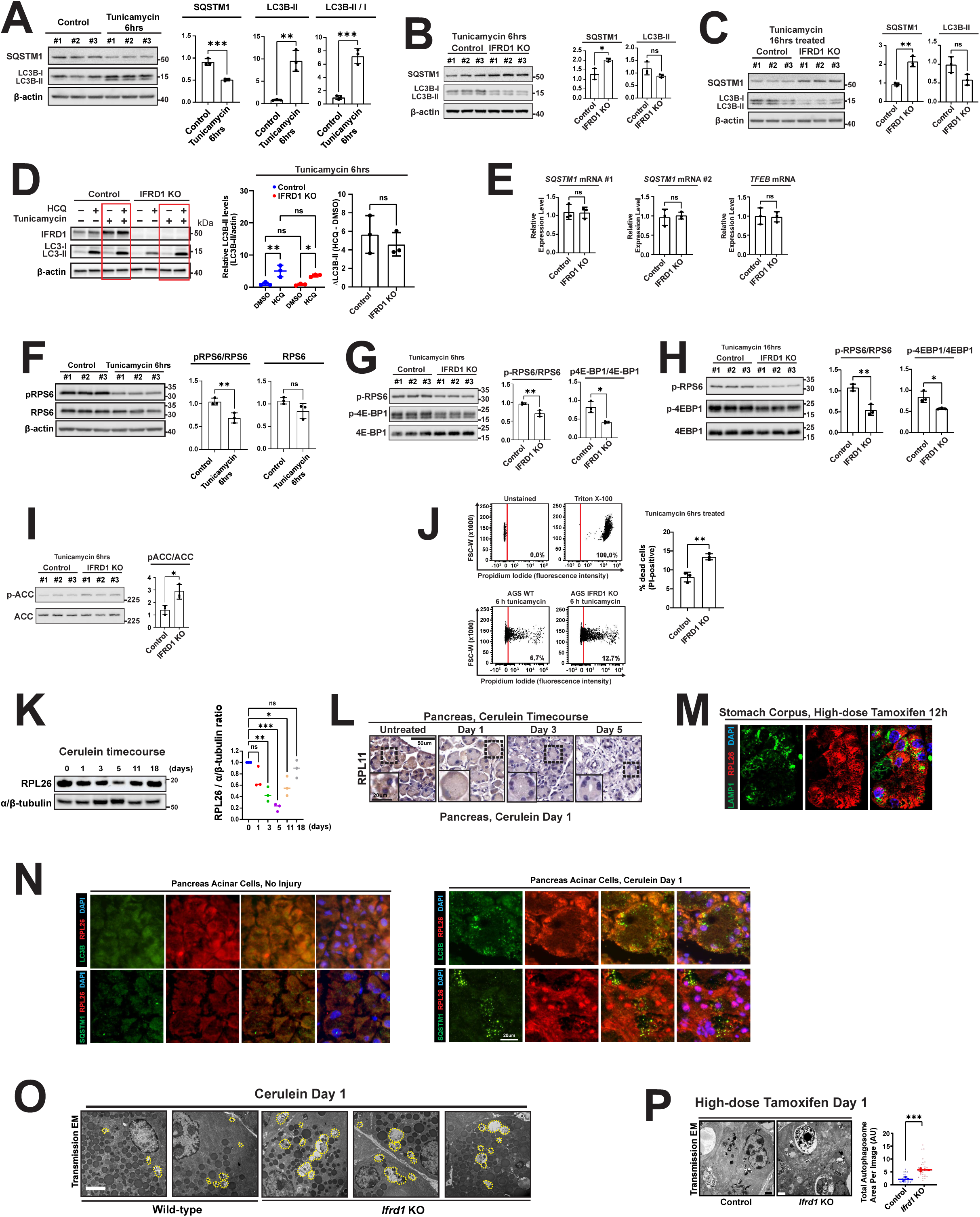

We next investigated whether the defect in ribosome salvaging caused by loss of IFRD1 leads to secondary metabolic derangement. Tunicamycin-induced injury is known to suppress mTORC1 activity, as manifested by decreased phosphorylation of RPS6 (Figure 6F). We hypothesized that cells lacking IFRD1 would suffer an even more pronounced reduction in mTORC1 activity, consistent with our previous observations during paligenosis in the stomach and pancreas of IFRD1-knockout mice^12^. Indeed, we observed that IFRD1-knockout cells exhibited further significant reductions in the phosphorylation of mTORC1 targets. Specifically, lower pRPS6/RPS6 and p4E-BP1/total 4E-BP1 ratios were detected within 6 hours of tunicamycin treatment (Figure 6G) and persisted through 16 hours of treatment (Figure 6H), confirming impaired mTORC1 function in the absence of IFRD1.

Because AMPK is a primary responder to perturbations in energy homeostasis and a known inhibitor of mTORC1, we reasoned that AMPK activation might drive this further reduction. Supporting this, we detected an increase in the pACC/ACC ratio (Figure 6I), a widely established downstream marker of AMPK activity^38,39^.

Collectively, these data suggest that perturbed ribosomal regulation in the absence of IFRD1 results in a dual insult: the overburdening of the autophagic machinery and AMPK-mediated oversuppression of mTORC1. This metabolic collapse ultimately drives increased cell death (Figure 6J), as demonstrated by an increased proportion of propidium iodide-positive cells in the IFRD1 knockout group in flow cytometric analysis of cell viability.

We finally investigated ribosomal dynamics in vivo in paligenosis during which IFRD1 plays a crucial role to stabilize cells to eventually induce mTORC1 after the initial massive upregulation of autophagy in stage 1. We noted that, consistent with ribophagy being a critical aspect of the autophagy in paligenosis, there was a dramatic reduction in ribosomal proteins, such as RPL26 after cerulein treatment (Figure 6K), which was further confirmed by the loss of RPL11 signal intensity in the acinar cells undergoing paligenosis (Figure 6L). The autophagic degradation of ribosomes was corroborated by overlap between lysosome/autolysosome marker LAMP1 and RPL26 in chief cells in the stomach after both high-dose tamoxifen-induced gastric injury (Figure 6M) and in acinar cells during pancreatitis (Figure 6N).

We hypothesized, given IFRD1’s role in ribosomal salvage and in clearance of p62-tagged substrates during autophagy that we saw in vitro, that *Ifrd1* null cells would show similar defects in autophagy. Accordingly, transmission electron microscopy (TEM) revealed that pancreatic acinar cells from global *Ifrd1*-knockout mice displayed larger and more numerous autophagosomes and autolysosomes (Figure 6O). Similarly, gastric chief cells lacking IFRD1 exhibited significantly larger autolysosomes following high-dose tamoxifen treatment compared to wild-type controls (Figure 6P; 5.80 ±0.51 vs. 2.25 ±0.74 µm², *P* = 0.0006).

In summary, these data demonstrate that the absence of IFRD1 and the failure to salvage ribosomes leads to altered autophagic flux with increased delivery of non-salvaged ribosomes into the autophagic pathway, and reduced mTORC1 activation or reactivation.

## Discussion

In this study, we demonstrate that IFRD1 functions as a bona fide ribosome-binding protein that serves as a ribosome salvaging factor. This ribosome salvaging mechanism adds to our understanding of how global translation can be regulated, beyond phosphorylated eIF2α’s well-characterized role in inducing global translational arrest and SG formation. IFRD1 serves as a distinct, independent "caretaker" of the ribosome during cell-stress-induced translational blockade.

Although IFRD1 would be expected to primarily stabilize only the ribosomes it binds, IFRD1 loss during stress causes more global cellular shifts in mTOR and autophagy. The results here explain why IFRD1’s role in preserving ribosomes under stress and in transition, would be important in cellular regeneration programs like paligenosis wherein stress causes cells with abundant ribosomes (largely on rough ER) to downscale into proliferating cells that need cytoplasmic ribosomes to replicate cellular components for cell division. We also note that the results here paint a much fuller picture of IFRD1’s role in vivo during paligenosis. We show here and previously that loss of IFRD1 in *Ifrd1-/-* mice during paligenosis in vivo and in tunicamycin-induced stress in vitro both lead to aberrant autophagy with subsequent failure to maintain and/or reactivate mTORC1^12^.

IFRD1 joins an emerging repertoire of ribosome-binding proteins that can maintain a pool of idling (i.e., non-translating) ribosomes, which challenge the dogmatic view that assembled 80S ribosomes are largely not idle, spending all their lifespan actively translating protein. In prokaryotes, factors such as Hibernation Promoting Factor (HPF) and Ribosome Modulation Factor (RMF) induce the dimerization of 70S ribosomes into inactive 100S complexes^40-43^, as well as universal stress protein (USP) that can maintain nonreplicative persistent state^44^. In yeast, Stm1 (the mammalian SERBP1 homolog) binds and clamps ribosomes into an 80S state^45-47^. In multicellular eukaryotes, SERBP1 and IFRD2 act as molecular clamps, threading through mRNA entry/exit channels and the A and P sites to prevent translation^29,48^. These ribosome-binding proteins vary significantly in their tissue expression, triggers, and behaviors. For example, while eIF2α phosphorylation induces IFRD1, SERBP1 is inhibited and phosphorylated via mTOR^48^. IFRD2 is expressed in rabbit reticulocytes (where IFRD1 is absent), whereas Drosophila gonads show sex-specific expression of IFRD1^29,30^, in both systems, even in the absence of acute injury or stress. Furthermore, the consequences of losing these factors differ. In our study, IFRD1 loss resulted in a more pronounced reduction of the 60S peak relative to the 40S and 80S peaks, suggesting favored 60S-specific degradation. A plausible explanation for the IFRD1 knockout phenotype is that 40S subunits are prioritized for preservation, as they more readily re-engage in subsequent translation or localized to SGs^49,50^. Interestingly, reduced 80S ribosomes were accompanied by an increase in free 40S and 60S subunits in *Serbp1* knockouts, suggesting that the specific stress context or the specific binding protein may dictate the pattern of ribosomal decay; however, both subunits seem to be ultimately degraded as noted in the later timepoints in *Ifrd1* knockouts.

Interestingly, among the ribosomal proteins examined with and without IFRD1, only RPL11 showed a significant reduction without IFRD1, potentially reflecting that different proteins of 60S subunit have differing half-lives, with some more readily degraded via the ubiquitin-proteasome pathway^35^. In paligenosis, IFRD1 loss leads to p53-mediated death, which can be caused by accumulation of specific large ribosomal subunits like RPL11^51^, and not small ones. That would be another reason why the preferential depletion of 60S units in the absence of IFRD1 is so damaging, and also an indication for why IFRD1 is important for paligenosis. In our in vitro studies, links between ribosomal degradation and p53 were difficult to establish, likely due to an aberrant p53 regulation frequently noted in cancer-dervied cell lines, even with wildtype p53^52,53^.

While IFRD1 clearly is found on non-translating ribosomes, how and when IFRD1 associates with those ribosomes remains unclear. We propose two theories: first, IFRD1 may occupy ribosomes immediately following, or concomitant to, translation termination, thus preceding (or outcompeting) recycling factors like ABCE1^54,55^; or second, IFRD1 could associate with the 5’ UTR or initiator tRNA during early translation initiation. Furthermore, the post-injury context, specifically how translation resumes, also remains poorly understood. In yeast, the Dom34-Hbs1 complex (orthologous to human PELOTA-HBS1L) facilitates the deactivation of Stm1, "declamping" ribosomes to return them to the active pool^56^. Whether a similar mechanism exists for IFRD1 is a key question for future research. PELOTA-HBS1L or ABCE1 might actively displace IFRD1, or IFRD1 may simply undergo spontaneous dissociation to bind another ribosome as described above.

These findings open new avenues for investigation. In particular, our studies mostly use a non-paligenosis, in vitro model of stress to induce IFRD1 and determine its core, evolutionarily conserved mechanism of action, but we speculate the results have broad implications for the inherent logic of paligenosis. It has been shown that differentiated cells are primarily enriched in rough ER ribosomes because differentiation is largely about increasing production of specialized cellular function, most of which involve interaction with other cells or modification of extracellular space. Thus, differentiated cells need secreted or membrane-associated proteins, whereas stem cells largely care about internal proteins: making copies of ribosomes and histones and housekeeping genes for cell replication^57^. For cells to undergo paligenosis from differentiated to stem-like state, they must have a way of preserving and relocating their ribosome pool. IFRD1 may be necessary for shepherding ribosomes safely through key transitions like loss association to mRNA and/or movement of ribosome from rER to cytosol. Perhaps, once the ribosome navigates the transition, may not need to remain bound to a ribosome to keep it stabilized, such that a single IFRD1 molecule can stabilize multiple ribosomes.

One view for the conserved nature of paligenosis is that it is an ancient program for multicellular organisms to allow maximal differentiation and specialization of cell types while preserving the ability to recruit those cells as stem cells when there is tissue injury^58^. That ability comes with a price as differentiated cells may have accumulated abnormal genomes that put them at risk for aberrant clonal expansion if they regain proliferative potential. Hence, p53 evolved with paligenosis to check cellular health before allowing cells transitioning from stage 2 to stage 3 to complete paligenosis. Our results show how ribosomal salvage and translocation can potentially be monitored by p53 in the stage 3 paligenosis checkpoint.

Similarly, a recent study revealed mechanisms underlying the stage 1 to 2 checkpoint. In stage 1, cells undergo massive autophagy to downscale mature architecture before proceeding to Stage 2, when then begin to express progenitor and proliferation-related genes. It was shown that progression to stages 2 and 3, depends on induction of the Hippo signaling effector YAP1^59^. YAP1 induction, in turn, depends on the autophagy of stage 1, because STK38, a kinase that constitutively targets YAP1 for degradation is degraded by autophagy. Thus, transition to Stage 2 depends on autophagy in Stage 1. Given the evolutionary conservation of IFRD1 and the Hippo pathway proteins as stress-induced, and regeneration-promoting, we are gaining some insights into potential universal cellular logic in paligenosis. However, of course, much more work across many tissues and injury patterns must be done to provide more evidence for this theory.

## Methods

### Mice

All experiments involving animals were performed according to protocols approved by the Baylor College of Medicine and Emory University Animal Studies Committee. *Nat10* conditional knockout mice were generated using CRISPR/Cas9 technology as previously published^23^. *Ifrd1* conditional knockout mouse was shared by RIKEN, and the specifics of the mice have been previously reported^20^. *Ifrd1* global knockout mice were maintained as described previously^12^. Tamoxifen (Toronto Research Chemicals, Inc, Toronto, Canada) was prepared in a 10% ethanol and 90% sunflower oil solution by sonication as previously described. Medium-dose tamoxifen (3 mg / 20 g mouse body weight, intraperitoneal injections) was administered for three consecutive days to induce recombination of Cre in the pancreas in *Mist1^CreERT2^* mice. High-dose tamoxifen was used to induce paligenosis in the stomach corpus of wild-type and *Ifrd1* global knockout mice following previously published protocols^60,61^. Cerulein (Bachem) was treated once at a concentration of 2 μg / 20 g for short term assay, and for six hourly injections, every other day, as previously described^62^. Puromycin was treated at the concentration of 50 μg / 20 g for 45 minutes before euthanasia.

### Cell culture and drug treatment

AGS cells were cultured in RPMI-1640 medium with 10% fetal bovine serum (FBS) and Primocin (100 µg/mL) or penicillin/streptomycin (100X). HEK-293T and LS-174T cells were cultured in DMEM medium with 10% FBS and Primocin (100 µg/mL) or penicillin/streptomycin (100X). All cells were maintained at 37°C in a 5% CO_2_ humidified atmosphere. Stable knockdown of IFRD1 was carried out using shRNA against IFRD1. IFRD1 knockout AGS cells were generated using CRISPR system. Tunicamycin was used at a concentration of 4 μg/mL from 4 to 24 hours of treatment. DMSO (≤ 0.1%) was used as controls. MG-132 was treated at a concentration of 10 µM for up to 5 hours. Sodium arsenite (NaAsO_2_) was used for treatment at a concentration of 0.25 mM (250 μM) for 30 minutes and thapsigargin was used at 10 µM concentration for 45 minutes. Puromycin was used for treatment at 5 μg/mL concentration for 15 minutes and replaced with regular media for 5 minutes. As a negative control, 50 mg/mL of cycloheximide was used 5 minutes prior to puromycin treatment. Live imaging of propidium iodide (1 μg/mL) and Hoechst 33342 (1 μg/mL) was carried out using previous methods.

### Plasmids

IFRD1 lacking amino acid residues 1-69 in the N-terminus (ΔIFRD1) was cloned in the pcDNA3.1 plasmid with an HA tag on the C-terminus, with help of Dr. Oskar Laur at the Custom Cloning Core Division, Emory Integrated Genomics Core, Emory University. (N)Td-Tomato-IFRD1, (C) IFRD1- HA- GFP, IFRD1- HA, Human IFRD1 with a N-terminal 6x-His tag, and transposase are gift from Dr. Jeffrey W. Brown at Washington University in St. Louis. Human IFRD1 with a N-terminal 6x-His tag was cloned into a pET-28a vector. Sequence was confirmed with sanger sequencing (genewiz). The vector was transformed into *Escherichia coli* BL21(DE3) and expression was induced with IPTG for 18 hours at room temperature. The cells were centrifugally collected and the pellet lysed in the presence of lysozyme and DNAseI with sonication. The centrifugally cleared supernatant was applied to an Ni-NTA column and eluted with a stepwise gradient 10-250 mM imidazole. The desired fraction (80 mM) was denatured in 6M GuHCl, 100 mM NaCl, 50 mM Tris pH 7 and then refolded by dialyzing against >100 volume equivalents of 500 mM Arginine, 100 mM NaCl, 1 mM DTT, 10 mM Tris pH 7 for 2 days. After which the chaototropic arginine was removed by dialysis against phosphate buffered saline and the protein in phosphate buffered saline was concentrated with Centricon centrifuge device.

### Western blot, immunoprecipitation, subcellular/ ER fractionation, and mass spectrometry data analysis for IFRD1, IFRD1-GFP Immunoprecipitation experiments

Western blotting was carried out in using the previously published protocol^23^. When cultured cells were used, the cells were trypsinized, and the pellets were washed in ice-cold PBS and centrifuged three times for 5 minutes at 4°C before lysis buffer was added. Subcellular fractionation was performed following the published protocol. For immunoprecipitation, we used a protocol previously described^23^. Briefly, lysates were prepared using Pierce IP lysis buffer (Thermo) and were incubated with 50 μL of Protein A beads with either an antibody of interest or normal rabbit IgG (negative control; Cell Signaling). To pull-down HA-GFP-tagged IFRD1, we used ChromoTek GFP-Trap® Magnetic Particles M-270 (Proteintech).

Spectral counts for IFRD1 3.5 hour and 20 hour samples after isolation by immunoprecipitation (IP) with anti-IFRD1 antibody were compiled for each protein identified. The same experiment using an immune-naive IgG was employed as a control and equivalent spectral counts for each identified protein were compiled. These spectral counts data for IFRD1-IP-3.5h, IgG-IP-3.5h, IFRD1-IP-20h, and IgG-IP-20h were converted to Normalized Spectral Abundance Factors (NSAF) to correct for total spectral counts for each sample and normalize for different protein molecular weights. Final correction to 10^6^ Total NSAF was used for final analysis. The NSAF difference between authentic IFRD1-IP versus Control-IP was calculated based on 10^6^ NSAF and used to generate a ranked protein signature. The IFRD1-GFP versus GFP control was isolated and subjected to untargeted proteomics analysis at the Baylor College of Medicine Proteomics Core Facility. The output spectral counts were processed using an in-house informatics pipeline to generate normalized and batch corrected data (represented as iBAQ values). These reported iBAQ data were used for comparative differential iBAQ analysis without further processing. Data from IFRD1 IP experiments was evaluated based on a differential NSAF value cutoff of +2.0. The positive differential NSAF indicates increased protein observed in the IFRD1 IP versus IgG control IP. The IFRD1-GFP data was evaluated for the top 500 proteins identified in the differential iBAQ signature. Proteins identified were submitted to the PANTHER (https://pantherdb.org) statistical overrepresentation test. Analysis parameters were PANTHER version 19.0, released 2024-06-20; Analyzed and Reference Lists were *Homo sapiens*; Annoation Data Set was PANTHER Protein Class; Fisher’s Exact Test with False Discovery Rate calculation. Data output was compiled in Python Version 3.11 for graphical analysis using Matplotlib (3.10.3), Matplotlib-venn (0.11.9) and Seaborn (0.13.2) libraries.

### Rabbit reticulocyte In Vitro Translation Assay

Rabbit reticulocyte luciferase mRNA translation assays were performed according to the manufacturer’s protocol (Promega). Briefly, 35 µL of nuclease-treated rabbit reticulocyte lysate was incubated with water, bovine serum albumin (3 mg/mL), or various concentrations of IFRD1 purified from *E. coli*, along with luciferase control RNA (1 µg/µL, 2 µL) and an amino acid mixture to a final volume of 50 µL. The reactions were incubated for up to 1 hour at 30°C in a heat block, and luminescence was measured using a luminometer (BioTek).

### ER fractionation

AGS cells were seeded in 10 cm dishes to reach ∼80–90% confluence on the day of the experiment. Cells were treated with tunicamycin (8 μg/mL) for 2, 4, or 6 hours prior to harvest. Before fractionation, cells were incubated with 10 mL of ice-cold PBS containing 50 μg/mL cycloheximide (CHX) for 10 minutes on ice to stabilize ribosome-associated complexes. For cytosolic extraction, 1 mL of permeabilization buffer (110 mM KOAc, 25 mM K-HEPES pH 7.2, 2.5 mM Mg(OAc)₂, 1 mM EGTA, 0.015% digitonin, 1 mM DTT, 50 μg/mL CHX, 1× Complete Protease Inhibitor Cocktail, and 40 U/mL RNaseOUT™) was added to each dish. Cells were incubated at 4°C on a rotating platform for 5 minutes. The released soluble material was collected as the cytosolic fraction. Cells were then gently washed with 5 mL of wash buffer (110 mM KOAc, 25 mM K-HEPES pH 7.2, 2.5 mM Mg(OAc)₂, 1 mM EGTA, 0.004% digitonin, 1 mM DTT, 50 μg/mL CHX). For ER-enriched membrane extraction, cells were incubated with 1 mL of lysis buffer (400 mM KOAc, 25 mM K-HEPES pH 7.2, 15 mM Mg(OAc)₂, 1% NP-40, 0.5% sodium deoxycholate (DOC), 1 mM DTT, 50 μg/mL CHX, 1× Complete Protease Inhibitor Cocktail, and 40 U/mL RNaseOUT™) at 4°C on a rotating platform for 5 minutes. The NP-40–soluble material was collected as the ER-enriched fraction. Both cytosolic and ER fractions were clarified by centrifugation at 13,000 rpm for 10 minutes at 4°C to remove insoluble debris. Supernatants were transferred to fresh pre-chilled microcentrifuge tubes and used for downstream western blot analysis.

### Imaging and tissue analysis

All cell and tissue IF and tissue IHC experiments were carried out using a previously published protocol^61^. Cultured cells were grown on coverslips in 12-well culture plates or 4-well chambers. IF images were captured using the AX R Confocal System with Eclipse Ti2-E Inverted Microscope (Nikon), LSM880 Laser Scanning Confocal Microscope (Zeiss), and Leica DM6B upright fluorescent microscope, ensuring consistent exposure times across samples for comparison. X-gal staining was done as previously described^10^.

### Sucrose density gradient centrifugation

Sucrose density gradient centrifugation was carried out according to a manual fraction protocol^23^, or using piston gradient fractionator (BioComp) to measure a continuous OD260 value using the TRIAX system to compare the gradient peaks in IFRD1 knockout samples vs. control samples. When conducting run-off assays, cycloheximide was omitted in all steps. To block the initiation step, harringtonine was used for treatment at a concentration of 2 μg/mL for 2 minutes. When UV crosslinking was used, 200 mJ/cm^2^, was applied once, without lid, in 3 mL ice-cold PBS, and then lysis buffer was added immediately. For formaldehyde crosslinking, cells were incubated on 5 mL of 0.4% PFA in PBS for 5 minutes, and washed in PBS before the lysis buffer was added.

### RNA extraction and qRT-PCR

RNA was isolated using the RNeasy Mini kit (Qiagen) or Direct-zol RNA Miniprep kit (Zymo Research), and also underwent on-column DNase digestion per the manufacturer’s protocol. The quality of the mRNA was verified with a Nanodrop spectrophotometer (Thermo). 500 ng of RNA was reverse transcribed with the PrimeScript^TM^ RT Reagent kit (Takara) following the manufacturer protocol. Power SYBR Green Master Mix (Thermo) fluorescence was used to quantify the relative amplicon amounts of each gene (see Supplementary Table 3 for the list of the primers). For the northern blot and RNA gel electrophoresis, we extracted RNA from the pancreas using TRIzol^TM^ reagent using the previously used protocol^23^.

### Transmission electron microscopy

Pancreas preparation and imaging for transmission electron microscopy were performed as previously described^23^. Briefly, whole pancreas was collected, fixed overnight at 4°C in modified Karnovsky’s fixative and sectioned. Pancreas tissue was processed and imaged at the Molecular Microbiology Imaging Facility, Department of Molecular Microbiology, Washington University in St. Louis.

### Cell Viability Assay

IFRD1 wild-type and IFRD1 CRISPR knockout AGS cells were seeded at approximately 0.1 x 10^6^ – 0.3 x 10^6^ cells per well in 6-well cell culture plates and incubated at 37°C under 5% CO_2_. Upon reaching 100% confluency (48 – 72 h after seeding), cells were incubated in the absence or presence of tunicamycin (4 µg/mL, 0.08% DMSO). After 6 h, cells floating in culture media were collected into labeled microcentrifuge tubes and monolayers were detached from cell culture plates with TrypLE and combined with the corresponding detached cells. Cells were centrifuged at 300 x g, 5 min at 4°C to pellet, supernatants were discarded and pellets were resuspended in propidium iodide staining solution (1X PBS containing 10 µg/mL propidium iodide, 1% BSA, and 1 mM EDTA). Positive control cells were resuspended in propidium iodide staining solution containing added 0.1% Triton X-100 to permeabilize cell membranes. Negative control cells were resuspended in 1X PBS containing 1% BSA and 1 mM EDTA (unstained). Cells were incubated in staining solution on ice, protected from light. After 15 minutes, cells were analyzed for propidium iodide fluorescence by flow cytometry with a BD FACS Canto II analyzer using a 561 nm excitation laser, 600 nm long-pass emission filter, and 610/20 nm bandpass emission filter. Single cells were identified using forward scatter (FSC) and side scatter (SSC). BDFACS FSC 3.0 files were analyzed using FlowJo 10 software. The 2D dot-plots are representative of three biologically independent replicates (n = 3). The data from three biologically independent replicates (n = 3) were combined and the percentage of dead cells per condition was calculated using min-max normalization with the mean of unstained cells representing the minimum (0.0) and the mean of Triton X-100 permeabilized cells representing the maximum (100.0). Data are presented as mean ± SD with individual data points representing the value from an individual replicate.

### Statistical analyses

Statistical analyses were performed using Prism 10 (GraphPad). In general, if datapoints were normally distributed, a Student’s t-test (for 2-sample assessments of error) was used to quantify the likelihood of true differences in means. For paired samples, a paired t-test was used. Where distribution was non-normal, nonparametric tests were used. For comparisons between multiple groups, a one-way ANOVA with Dunnett’s post hoc test or a two-way ANOVA with a multiple comparisons test was used to determine significance. *P* < 0.05 was considered statistically significant for interpretation in the text.

## Supporting information

Graphical Abstract

## Abbreviations

IP: Immunoprecipitation
SG: Stress Granule
IF: Immunofluorescence
IDR: Intrinsically disordered region
ER: Endoplasmic reticulum
RRL: Rabbit reticulocyte lysate
RiBi: Ribosome biogenesis

## Author contribution

CJC: experiment design, supervision of the experiments, data acquisition, analysis, interpretation, funding, writing of the manuscript; MKC : experiment design, acquisition, analysis, interpretation, writing of the manuscript, AKR: experiment design, data acquisition, analysis, interpretation, TN: experiment design, data acquisition, analysis, interpretation, writing of the manuscript; SJB: bioinformatic analysis, interpretation, writing of the manuscript, SL: experiment design, data acquisition, analysis, interpretation, JWB: experiment design, acquisition, analysis, interpretation, writing of the manuscript, JCM: experimental design, supervision of the experiments, data interpretation, funding, writing of the manuscript. All authors reviewed the manuscript.

## Data and code availability

The mass spectrometry proteomics data have been deposited to the ProteomeXchange Consortium via the PRIDE partner repository with the dataset identifier PXD077756.

## Acknowledgments

We thank the RIKEN BDR animal facility for providing the *Ifrd1* floxed mice (Accession No. CDB0833K). The Baylor College of Medicine Mass Spectrometry Proteomics Core is supported by the Dan L. Duncan Comprehensive Cancer Center Award (NIH/NCI, P30CA125123) and Core Facility Award from the Cancer Prevention and Research Institute of Texas (RP210227). The expert technical assistance of Petra Erdmann-Gilmore, Dr. Yiling Mi, Alan Davis and Rose Connors is gratefully acknowledged. The proteomic experiments were performed at the Washington University Proteomics Shared Resource (WU-PSR), R Reid Townsend MD.PhD., Director and Robert Sprung, PhD., Co-Director). The WU-PSR is supported in part by the WU Institute of Clinical and Translational Sciences (NCATS UL1 TR000448), the Mass Spectrometry Research Resource (NIGMS P41 GM103422; R24GM136766) and the Siteman Comprehensive Cancer Center Support Grant (NCI P30 CA091842). Some of the microscopy and all tissue sectioning were performed by the Tissue Analysis & Imaging Core of the Washington University Digestive Disease Research Core Center (NIH/NIDDK, P30DK052574) or by the Tissue Analysis and Molecular Imaging (TAMI) Core of the Texas Medical Center Digestive Disease Center (NIH/NIDDK, P30DK056338). This project was supported by the Cytometry and Cell Sorting Core at Baylor College of Medicine with funding from the CPRIT Core Facility Support Award (CPRIT-RP240432), the NIH (CA125123 and ODO36336), and the assistance of Joel M. Sederstrom. We also thank Robert Lawrence, Ph.D., E.L.S. for editorial support in review of the manuscript.

