## Supplementary Table 1 for "Stress-Responsive Protein IFRD1 Protects Assembled Ribosomes via a Ribosome-Salvaging Mechanism"

| IFRD1_IP_3 | IFRD1_IP_2 | GFP_IP |
|---|---|---|
| YBX1 | YBX1 | IFRD1 |
| RPL26L1 | NONO | RPLP2 |
| RPL31 | SFPQ | RPL29 |
| G3BP1 | G3BP1 | NPM1 |
| RBMXL1 | PABPC3 | RPS11 |
| RPS10 | PABPC1 | RPS3A |
| RPL26 | RPS15A | RPL3 |
| SRSF1 | HNRNPA2E | RPL14 |
| H2BC3 | RPS10 | RPL13 |
| RPS9 | SRSF1 | RPL5 |
| SRSF3 | DDX6 | RPL12 |
| RPL15 | RPL26L1 | BCLAF1 |
| DYNLL1 | ERH | RPL10A |
| RPS21 | RBMXL1 | NCL |
| H1-4 | RPS24 | RPL26 |
| RPL30 | RBMX | RPL7A |
| DDX6 | HNRNPK | RPLP0 |
| NUDT21 | SRSF3 | RPL35 |
| RPL37A | IFRD1 | RPS6 |
| SART1 | HNRNPA3 | RPL23A |
| RPS24 | H2BC3 | RPS28 |
| PABPC1 | RPL31 | RPL8 |
| SLC25A6 | TIMM13 | THRAP3 |
| FUBP3 | UPF1 | RPS15 |
| RPL28 | NUFIP2 | RRBP1 |
| TIMM8B | ELAVL1 | RPS23 |
| RPS2 | FXR2 | RPL4 |
| RELCH | MAGOH | HIST1H1C |
| ACTC1 | RPL15 | RPS14 |
| H1-10 | SRSF9 | LACTB |
| H1-3 | ACTC1 | RPL28 |
| RPL7A | RPL32 | HNRNPM |
| RBM47 | RELCH | RPL32 |
| MSI2 | G3BP2 | PLXNA4 |
| DDX3X | RPS27 | RPL35A |
| RPL13A | RPL36A | RPL34 |
| RPL9P8; RI | MOV10 | RPL13A |
| RPL32 | HNRNPH1 | RPL21 |
| MYCBP | DDX3X | RPL17 |
| RPL36A | RPS8 | RPL23 |
| RACK1 | DDX5 | RPL11 |
| PABPC1L | H1-4 | RPS4X |
| TUBA1B | HNRNPM | RPL19 |

| | | |
|---|---|---|
| RPL34 | LARP1 | RPL7 |
| HNRNPH1 | RPS13 | HIST1H1B |
| KHSRP | RPS20 | RPS8 |
| BCLAF1 | RBM8A | DDX3X |
| RPL3 | FUBP3 | RPL17-C18orf32 |
| LARP4 | RPS14 | RPS10 |
| RPS29 | RPL24 | RPL22 |
| EIF4G1 | PABPC4 | RPS13 |
| SRSF9 | FAM120A | RPL27 |
| HNRNPL | EIF2S1 | RPS7 |
| LARP4B | FAM98A | RPL36AL |
| HMGA1 | RTCB | RPS25 |
| ILF3 | RPS23 | RPL27A |
| ZCCHC3 | TIMM8B | RPL31 |
| CEP55 | SART1 | RPS27 |
| RPS15A | IGF2BP3 | YBX1 |
| DYNLL2 | GDF15 | RPL6 |
| H1-5 | MATR3 | RPS2 |
| WDR13 | EIF2S2 | TOP1 |
| GATA6 | HNRNPA1 | RPL26L1 |
| TRA2B | HP1BP3 | RPL15 |
| HNRNPH3 | YBX2 | RPL18A |
| MATR3 | MAGOHB | RPL36A-HNRNPH2 |
| SRSF7 | WDR13 | RPL36A |
| LARP1 | RTRAF | DDX5 |
| TIMM8A | MSI2 | RPL37A |
| LSM2 | ILF3 | RPS18 |
| EIF2S1 | FUS | H1F0 |
| HP1BP3 | C11orf98 | HNRNPUL1 |
| SNRPB | TUBA1B | DDX17 |
| DDX1 | DDX1 | SRP14 |
| HNRNPA3 | RPL9P8 | RPL10 |
| TDRD3 | ATXN2L | RPL24 |
| C11orf98 | RPS2 | RPL30 |
| RPS15 | SERBP1 | SRSF1 |
| RPL4 | RPL26 | RPL36 |
| RBM3 | EIF4E | RPS24 |
| SRSF6 | RACK1 | MTDH |
| KIF1C | STAU2 | RPS5 |
| DAZAP1 | FMR1 | DHX9 |
| PTPN3 | HNRNPL | H1FX |
| EIF3E | RPL5 | SPATS2L |
| HNRNPA2E | PSPC1 | RPL9 |
| IFRD1 | DHX30 | ATAD3B |

| | | |
|---|---|---|
| RPLP1 | ALYREF | SRP9 |
| SNRPGP15 | KHSRP | KRT37 |
| LSM6 | CAPRIN1 | PLA2G2A |
| STRBP | BCLAF1 | SSB |
| RPL10 | RPL30 | HNRNPU |
| RPS17 | RPS5 | RPS9 |
| RPL21 | RBM47 | SRSF5 |
| YBX2 | H1-5 | TBL2 |
| HNRNPH2 | USP10 | HSPD1 |
| PCBP3 | SRSF7 | ADAR |
| SRP9 | DDX17 | ILF3 |
| ATXN2 | SRSF6 | PNN |
| CAPRIN1 | SHFL | UBTF |
| RPL27A | EIF3E | ACTG2 |
| MARK2 | RPL18A | SNRPD2 |
| CEP44 | IGF2BP2 | APOBEC3C |
| MTCL1 | HNRNPUL1 | ZC3HAV1 |
| HNRNPA0 | LARP4 | ACTA2 |
| PFDN2 | HMGA1 | CAPRIN1 |
| KHDRBS1 | EIF2S3 | RBMX |
| EPS15 | RPS29 | MKI67 |
| USP10 | ZCCHC3 | SRSF7 |
| SNRPD2 | HNRNPF | ERH |
| SNRPD3 | TDRD3 | SFPQ |
| HNRNPF | EIF3A | LYAR |
| EIF4E | SLC25A6 | RACGAP1 |
| EIF4A2 | STRBP | PTBP3 |
| PRRC2C | AGO2 | EZR |
| CCDC9B | CEP44 | GNL3 |
| AGO3 | SRP9 | MOV10 |
| TIMM13 | EIF4A3 | NEBL |
| DDX50 | LARP4B | RPL18 |
| ELAVL2 | EIF3C | ZC3H18 |
| SRPK1 | RPL23 | C11orf98 |
| HBA1; HBA | GATA6 | SRPK1 |
| DDX21 | TIMM8A | BAG2 |
| PPP1CA | SYNCRIP | YTHDF1 |
| PKP2 | HNRNPH3 | FAU |
| YTHDF2 | PKP2 | LRRC59 |
| MOV10 | PTPN3 | YME1L1 |
| STAU2 | HNRNPU | PRPF19 |
| ZC2HC1A | CEP55 | CALML5 |
| HNRNPU | NCBP2 | KHDRBS1 |
| CPSF6 | ATP5F1C | MRPL11 |

| | | |
|---|---|---|
| DHX15 | SF3B6 | ALYREF |
| RPL36 | LSM6 | UHRF2 |
| FAM120A | SLC25A4 | RBM28 |
| RPAP3 | YBX3 | GRWD1 |
| RSBN1 | UPF3B | TRMT1L |
| FMR1 | NCBP1 | CKAP4 |
| SHFL | RPL7A | RBM6 |
| UPF2 | RPAP3 | SAP18 |
| PRPF6 | CIRBP | TBCEL |
| HNRNPR | ATXN2 | TRA2B |
| SPATS2L | EIF4G1 | CDX2 |
| HNRNPUL1 | TNRC6B | PPIL2 |
| POLR2E | PCBP3 | DDX24 |
| POLDIP3 | RPL21 | SERBP1 |
| RPL5 | ATP5PO | RBM10 |
| RPS20 | PRRC2C | FXR2 |
| ADD1 | EPS15 | LLPH |
| SSB | NUDT21 | AP2B1 |
| URI1 | EIF3G | ZC2HC1A |
| PURA | URI1 | NOP2 |
| SNRNP40 | DYNLL2 | RRP1B |
| CHTOP | DYNLL1 | FAM192A |
| RRBP1 | CASC3 | ASPH |
| YTHDF1 | DDX21 | HIST1H1D |
| NCBP1 | MTDH | KIF23 |
| SRSF4 | PFDN2 | SPATS2 |
| ASXL2 | RPL6 | MSN |
| PGAM5 | PTBP1 | NOP53 |
| ZC3H18 | AGO1 | POLDIP3 |
| SRSF10 | PRRC2A | DHX15 |
| RPS5 | SNRNP40 | DHX30 |
| STAU1 | RPS17 | NVL |
| RBM4B | RPS11 | RBM39 |
| RBM4 | RPL4 | ELOA |
| RBMX | UXT | CNTNAP4 |
| SERBP1 | PURA | PWP1 |
| RBM17 | EIF3H | TMCO1 |
| VAPB | YTHDF2 | EXOSC10 |
| BRK1 | RPS4X | KTN1 |
| EIF3A | LSM2 | MARK2 |
| EIF3C | HNRNPH2 | RBM14-RBM4 |
| PRRC2A | RPS9 | SRPK2 |
| RUVBL2 | RUVBL2 | HP1BP3 |
| UNK | RPL10 | EIF2S2 |

| | | |
|---|---|---|
| SNRPA1 | EIF3B | HNRNPA1 |
| TRA2A | HNRNPUL2 | MAP7 |
| TNRC6B | ZBTB2 | C7orf50 |
| RPS6 | STAU1 | RFC4 |
| EIF2S2 | C1QBP | EPRS |
| AKAP8 | AGO3 | SRP68 |
| MACO1 | SSB | PGAM5 |
| ZBTB2 | RUVBL1 | OASL |
| RPL7 | EIF3D | RNPS1 |
| FUS | LYAR | NAT10 |
| DHX36 | ELAVL2 | AP2A1 |
| AGO2 | RPLP1 | ZFP36L1 |
| HNRNPM | LSM3 | NIFK |
| GDF15 | TAF6 | GLUL |
| DHX30 | CCDC9B | YTHDF2 |
| CCDC9 | THRAP3 | CCDC59 |
| SF3B6 | ZCCHC8 | RIOX1 |
| LSM8 | SNRPB | AKAP8 |
| ZC3HAV1 | EIF3K | KARS |
| PABPC4 | TUBA8 | DDX52 |
| AGO1 | MRPL11 | NOL7 |
| SAFB2 | ZC3HAV1 | DDX27 |
| BLTP3B | PPP1CA | NOSTRIN |
| SNRPA | HNRNPA0 | DHX36 |
| UPF3B | RPL3 | CLASRP |
| CSDE1 | TRA2B | SRSF6 |
| H1-0 | HNRNPR | ECT2 |
| WDCP | EIF3I | RFC1 |
| RSBN1L | L1RE1 | GPATCH2 |
| PPP1CC | EIF3M | MRPS23 |
| HNRNPK | CHTOP | GTPBP4 |
| ALYREF | SNRPD1 | DKC1 |
| PAIP1 | EIF1B | ASZ1 |
| PTBP3 | MARK2 | MYBBP1A |
| SAFB | PRPF6 | PRC1 |
| TACC2 | RBM17 | MTPAP |
| AP2A2 | SNRPGP15 | SDAD1 |
| NCOA5 | NDUFA4 | RCC2 |
| ASDURF | SF3B2 | RBM17 |
| TAF9B | SRRT | SRP72 |
| ZCCHC8 | POLDIP3 | KRR1 |
| EIF2S3 | DDX50 | SRRM2 |
| PTPN13 | EIF3L | EBNA1BP2 |
| ZNF638 | SRPK1 | SRPRB |

| | | |
|---|---|---|
| SNRPC | PPP1CC | SYNCRIP |
| RPL19 | EPCAM | SIAH1 |
| DNAAF10 | SNRPD3 | TOP2A |
| EIF4A3 | SNU13 | CDC5L |
| BCKDK | TAF9B | YTHDF3 |
| CPSF7 | KIF1C | NOP16 |
| YTHDF3 | SPATS2L | FXR1 |
| TAF9 | CTTN | CD3EAP |
| CEP170 | DAZAP1 | DAZAP1 |
| OTUD4 | ZC3H18 | NKRF |
| LRRFIP1 | TAF9 | HMGCR |
| ZC3H11A | TOP3B | PRPF3 |
| SRRT | SLC25A1 | SEMA3B |
| PRPF8 | H1-10 | DDX18 |
| FAM98A | TOP1 | DDX55 |
| CPSF4 | PRKRA | EEF1E1 |
| RPL29 | RSBN1 | AFF4 |
| PHF5A | LARP1B | MARK3 |
| TOP1 | DNAAF10 | EXOSC3 |
| XRN1 | ZC2HC1A | TOP2B |
| TAF6 | RBM4 | CHD1 |
| RPL10A | LSM8 | TRIM25 |
| C1QBP | PTPN13 | CCDC25 |
| PPP1CB | ZC3H7A | RFC3 |
| ZC3H7A | KHDRBS1 | POLR1E |
| LARP1B | HNRNPD | DNAJB11 |
| SRSF5 | EIF4A2 | DNAJC9 |
| LYAR | MTCL1 | MOGS |
| SLAIN2 | CPSF7 | HADHA |
| STRAP | RPL35 | SF3B1 |
| EIF3D | EWSR1 | ZRANB2 |
| KHDRBS3 | ASXL2 | RRP15 |
| IGKV1-6 | DDX23 | SLAIN2 |
| EIF1B | DHX15 | SNRPA |
| RBM8A | RPL37A | DDX21 |
| EIF3B | H1-0 | GLYR1 |
| AP2B1 | ATP5MG | ASCC3 |
| ATP5F1C | TAF10 | YY1 |
| SF3B1 | SNRPE | SNU13 |
| EIF3H | C19orf53 | XRCC5 |
| YTHDC2 | ASDURF | RBM14 |
| RFC4 | CDKN2AIP | CKAP5 |
| PUM2 | YTHDF1 | KNOP1 |
| TAF8 | SNRPA1 | ZCCHC9 |

| | | |
|---|---|---|
| EIF4G3 | TRA2A | MLLT3 |
| PNN | NIFK | SLC16A1 |
| RPL18A | SLC25A11 | NUSAP1 |
| CTTN | SNRPC | TUBB3 |
| UPF1 | EIF3F | CMAS |
| CC2D1A | RBM4B | TRIP12 |
| YWHAZ | EFTUD2 | RARS |
| COX5B | ADAR | CBX4 |
| SNU13 | DHX36 | RFC2 |
| GNG5 | TACC2 | EIF4A1 |
| MARK3 | RBFOX2 | EIF2S1 |
| MRPL11 | DHX9 | AIMP2 |
| NOP16 | UPF2 | MELK |
| MTDH | HNRNPDL | STAU2 |
| RIOX1 | SAP18 | MAP4 |
| ADAR | MRPS26 | CHTOP |
| EIF3G | SUCLG1 | SRSF9 |
| SF3B2 | PHF5A | RBBP8NL |
| FAM83B | RPL39 | MRM3 |
| UBAP2L | RAB6A | IBTK |
| SUCLG1 | PRPF31 | OCIAD2 |
| TFAM | TAF15 | PPFIBP2 |
| RPL18 | SRSF10 | PCBP1 |
| TAF5 | POLR2E | PHF8 |
| ATP5PB | SRSF2 | SRRT |
| ZFR | ZNF638 | TJP2 |
| TRIM29 | RSBN1L | MRPS9 |
| TAF10 | AKAP8 | RBBP6 |
| MRPS23 | RFC5 | ACOT9 |
| EIF3I | PPIH | LARP1 |
| DDX23 | LRRFIP1 | EXOSC9 |
| RPL35 | PKP3 | EFCAB7 |
| SNRNP70 | YTHDF3 | SF3B6 |
| CELF1 | SAFB | CCDC9B |
| TOP3B | CEP170 | EIF2AK2 |
| AKAP1 | SRPK2 | NOP56 |
| KCTD14 | FGFBP1 | ZNF146 |
| GRSF1 | IGKV1-6 | RPP25 |
| NDE1 | YTHDC2 | ZNF281 |
| HNRNPUL2 | XRN1 | TRIM28 |
| RPL14 | PAIP1 | SF1 |
| APOBEC3B | UNK | DDX54 |
| RBM45 | BLTP3B | EPB41L5 |
| RPL13 | SURF6 | NKTR |

| | | |
|---|---|---|
| RBM7 | SNRNP200 | TCOF1 |
| PKP4 | CSDE1 | ABCF1 |
| RFC2 | RSL1D1 | HIST1H1E |
| YLPM1 | RRBP1 | HNRNPR |
| RPS23 | MACO1 | MRPS22 |
| HELZ | PTBP3 | CCDC77 |
| LUC7L2 | ILF2 | KIF2A |
| PRRC2B | MRPS6 | EXOSC5 |
| L1RE1 | PAM16 | FBXL13 |
| TAF4 | HELZ | EIF2S3 |
| TRIM25 | PRPF3 | MRPS7 |
| TFB1M | PRPF4 | TIGD6 |
| PIH1D1 | ATP5PB | UFM1 |
| MRPL54 | MRPS18B | MAGT1 |
| PDRG1 | ASPH | DDX60 |
| PLA2G2A | NFX1 | MATR3 |
| TOMM22 | ZFR | PAK1IP1 |
| NOP10 | NCOA5 | IFI16 |
| SUGP2 | CPSF6 | CLASP2 |
| SNRNP200 | CHERP | ARHGEF2 |
| RUVBL1 | SNRNP70 | GNL3L |
| EIF3L | FIP1L1 | SECISBP2 |
| AP2M1 | SUGP2 | MARS |
| TBL2 | KCTD14 | KIN |
| SRSF2 | LSM4 | REPIN1 |
| KDM1B | NDUFB4 | MAP7D1 |
| EFTUD2 | YLPM1 | AURKAIP1 |
| SPOUT1 | MKRN1 | BCAS2 |
| ZC3H7B | SAFB2 | VTN |
| SLC25A11 | PGAM5 | QARS |
| SZRD1 | PPP1CB | BUD13 |
| TXNL4A | WDR11 | LRRK2 |
| NONO | DHX29 | XRN2 |
| SPATS2 | SRSF5 | HDGFL2 |
| HNRNPLL | SF3B1 | YTHDC2 |
| CEP43 | PKP4 | CENPV |
| NELFE | CSNK1A1 | DHX57 |
| MRPS34 | KHDRBS3 | SLC29A3 |
| CDKN2AIP | WDCP | SPTY2D1 |
| LRPPRC | TAF5 | PAF1 |
| MARF1 | RFC4 | TBL3 |
| WTAP | MRPL54 | TDRD3 |
| GTF2IRD1 | PDRG1 | HS2ST1 |
| RRP1 | TOMM22 | SF3B3 |

| | | |
|---|---|---|
| ATP5PD | NOP10 | GPATCH4 |
| RNF41 | PRPF8 | CEP170 |
| TECR | WDR33 | RRS1 |
| DNAJB6 | PYM1 | MRTO4 |
| ZFP36L1 | PUM2 | RIOK2 |
| UXT | EIF5 | TUBA1A |
| UBL5 | RPL22 | MRPS18B |
| AP2A1 | SRP14 | MRPS35 |
| HNRNPD | SNRPD2 | EXOSC8 |
| GTPBP4 | MARK3 | SKP1 |
| BRI3BP | TAF4 | WDR36 |
| - | CPSF1 | SF3B4 |
| CHERP | CDX2 | TUBA1C |
| CPSF1 | RBM3 | EXD2 |
| RFC5 | NHP2 | HELZ2 |
| SKP1 | CYB5B | RBM27 |
| CIRBP | CPSF3 | MIS18BP1 |
| MAGT1 | CCDC9 | RRP7A |
| PPIH | SPATS2 | G3BP1 |
| DKC1 | RBM10 | PTCD3 |
| MRPS18B | DNAJA3 | UTP18 |
| PUM1 | MRPS34 | SRSF2 |
| PRPF3 | PRRC2B | PARS2 |
| SRPK2 | KRR1 | U2SURP |
| ADD3 | RRP1 | NOL6 |
| GRWD1 | AP2B1 | DARS |
| ZNF746 | ZC3H11A | GRSF1 |
| H1-2 | RPS3A | MAP4K4 |
| CIZ1 | SLTM | ERI1 |
| TAF7 | SLAIN2 | SDCBP |
| RBM42 | RNF41 | SCAPER |
| EN1 | DNAJB6 | ACIN1 |
| EXOSC8 | ZFP36L1 | SART1 |
| LSM5 | BRK1 | AATF |
| MED30 | TAF12 | NCBP1 |
| MBD6 | UBL5 | VWF |
| NCBP3 | MTREX | ELOB |
| HELZ2 | ZC3H7B | RSL1D1 |
| SURF6 | GMPPA | NFXL1 |
| RFC3 | ZBTB5 | NPM3 |
| CKAP5 | BRI3BP | SF3B2 |
| MAP7D1 | SEC13 | AKAP8L |
| SLBP | NSUN5 | CPT1A |
| RBM10 | PURB | MTREX |

| | | |
|---|---|---|
| DNAJA3 | CC2D1A | FARP2 |
| PRMT1 | ESRP1 | AHCYL1 |
| EPS15L1 | SNRNP27 | RBM34 |
| RRP1B | TARDBP | TNIK |
| LSM14B | RPS6 | ERCC3 |
| RPUSD4 | CCDC59 | EXOSC1 |
| MRPL12 | PUM1 | PARP1 |
| WDR11 | PCBP2 | PLRG1 |
| SECISBP2 | PRPF4B | RRP1 |
| MKRN1 | MBNL3 | CFAP20 |
| U2SURP | SF3A3 | IQCE |
| PURB | EIF5B | RBM15 |
| DGCR8 | AP2M1 | GPATCH8 |
| PABPN1 | TBL2 | RSBN1 |
| USP39 | CPSF4 | PES1 |
| CPSF2 | DNAJC9 | SURF6 |
| RSL1D1 | SRPRB | ESRP1 |
| CPSF3 | SF3B5 | KIAA1522 |
| RPL37 | LSM5 | IARS |
| PRPF31 | SF3A1 | LARP4 |
| RBMS2 | NCBP3 | RNF2 |
| PUSL1 | EIF4G2 | DNAJA2 |
| C7orf50 | TFAM | DIDO1 |
| SNRPE | EMD | KIF14 |
| ATP5MF | PNN | POLRMT |
| PACSIN2 | RFC3 | RACK1 |
| SLC25A1 | PHF6 | ELMSAN1 |
| CDX2 | RRS1 | BCL2L2-PABPN1 |
| NFX1 | RBM7 | SLTM |
| FXR2 | TRIM25 | QSOX1 |
| RNPS1 | CELF1 | DHX29 |
| SLC25A22 | FAM91A1 | LONP1 |
| EPCAM | UBAP2L | FAM207A |
| CCAR2 | MBNL1 | SART3 |
| BAP1 | RPS16 | KIF2C |
| GMPPA | AP2A2 | DIMT1 |
| KPNA2 | GRSF1 | ZCCHC8 |
| EBNA1BP2 | ZBTB44 | RO60 |
| NCKAP1 | PIH1D1 | SRPRA |
| MRM3 | MORF4L2 | DDX47 |
| CSNK1D | PFKP | CEP170B |
| DLG5 | NELFE | TTC3 |
| DECR1 | RBM45 | NOL10 |
| AXIN1 | RPSA | CCDC137 |

| | | |
|---|---|---|
| TSPAN10 | CPSF2 | KIF18A |
| RAB11A | WTAP | ERAL1 |
| LSM3 | GAR1 | ESF1 |
| TMA16 | MRPS23 | AIMP1 |
| RBBP7 | C7orf50 | ZNF638 |
| FAM120C | ATP5MF | ZNF512B |
| WDR33 | C17orf75 | TRIM56 |
| PFKL | YY1 | KRT23 |
| PFKP | RNPS1 | SNRPA1 |
| WDR5 | SLC25A22 | PARD3 |
| WWP2 | TAF8 | MRPS31 |
| GNL3 | HELZ2 | EXOSC7 |
| DDX18 | BLM | DEF6 |
| SEC22B | BAP1 | PSIP1 |
| THOC6 | BCKDK | ZNF568 |
| LUC7L3 | NUDT16L1 | ZNF598 |
| ZBTB44 | SSRP1 | RRP8 |
| IGKV1-27 | DKC1 | AP2M1 |
| FGFBP1 | DIMT1 | DDX60L |
| C1QC | EBNA1BP2 | ZNF622 |
| MBNL3 | RBM14 | UPF3B |
| RPL8 | U2SURP | MICU1 |
| ZNF326 | RRP1B | GTSE1 |
| DROSHA | AP2A1 | SGPL1 |
| ERI1 | TECR | RBM15B |
| EWSR1 | UQCRC2 | TAF10 |
| FAM91A1 | DDX28 | PCSK5 |
| CYFIP2 | TSPAN10 | MYO9B |
| RPS19 | MYCBP | TPX2 |
| RBFOX2 | DAD1 | REXO1 |
| HDAC1 | FAM120C | AURKA |
| ABI1 | AKAP1 | CHSY1 |
| KEAP1 | GRWD1 | EFTUD2 |
| TNRC6A | XRN2 | NUMA1 |
| TIAL1 | GNL3 | DNAJC10 |
| MBNL1 | DDX18 | ANKZF1 |
| F2 | SLX9 | EIF3D |
| SPOP | MRPL24 | RPN1 |
| SOX9 | TUBAL3 | EML3 |
| EXOSC6 | KLHL12 | CLASP1 |
| TIMM21 | MAGT1 | SNRPB2 |
| CDX1 | POLB | HNRNPUL2 |
| ATXN2L | SF3B3 | SUN2 |
| DHX29 | MARF1 | SETX |

| | | |
|---|---|---|
| YTHDC1 | DHX57 | ALDH3A2 |
| RPS14 | USP39 | CALU |
| BAIAP2L1 | RFC2 | PDCD6IP |
| CBX8 | IGKV1-27 | LARP7 |
| TIA1 | DCAF7 | SCO1 |
| PRPF4 | MRM1 | CDC20 |
| FRG1 | IGKV4-1 | CDK13 |
| CCDC59 | MOGS | TRAM1 |
| MRPL47 | PARP2 | LRBA |
| PHAX | TRIP4 | FMNL2 |
| TBC1D30 | AGR2 | MTHFSD |
| SF3A3 | CKAP5 | CBX8 |
| ZBTB5 | TAF7 | INAVA |
| CLK3 | TFB1M | SND1 |
| EXPH5 | RPS21 | TSR1 |
| SF3B3 | SF1 | ZC3H8 |
| KCNQ1 | EIF4G3 | |
| DDX28 | BRIX1 | |
| C17orf75 | NFIX | |
| YY1 | OTUD4 | |
| APOBEC3F | DDX55 | |
| PFDN6 | PACSIN2 | |
| LSM4 | COX5B | |
| BIN3 | PNO1 | |
| MRPS16 | EXOSC6 | |
| EXOSC3 | YTHDC1 | |
| PRSS33 | PNKP | |
| CYB5A | CIZ1 | |
| ZC3H13 | NCKAP1 | |
| SECISBP2L | YME1L1 | |
| XRN2 | CBX8 | |
| SLC25A10 | NSUN4 | |
| NKAP | SIN3A | |
| RPL12 | MUC6 | |
| NSUN5 | LRPPRC | |
| GSK3B | LACTB | |
| SYNCRIP | RBMS2 | |
| KLHL12 | PHAX | |
| PARP12 | CLK3 | |
| RNF213 | LAMP1 | |
| PRPF38B | PFDN6 | |
| HBE1 | EXOSC8 | |
| EIF1AX | MRPS16 | |
| SDCBP | EXOSC3 | |

| | |
|---|---|
| HBD | MRPS15 |
| SRSF8 | MRPS17 |
| RBM6 | SPTY2D1 |
| TRIM2 | CBX4 |
| MRPS2 | SLC25A10 |
| FIP1L1 | NKRF |
| BYSL | LUC7L2 |
| SLTM | KDM1B |
| PRPF4B | MRPS27 |
| SAP18 | EIF1AX |
| NIFK | CMSS1 |
| LSM14A | PLA2G2A |
| ACIN1 | RMI2 |
| EIF4G2 | SRSF8 |
| SLC3A2 | POP7 |
| SF1 | CHD6 |
| METTL17 | EPB41L5 |
| NHP2 | TRIM2 |
| FAM162A | MTPAP |
| NTHL1 | CCDC137 |
| POLR2H | PUSL1 |
| AP2S1 | GTPBP2 |
| MGST2 | BYSL |
| TAF3 | ST7 |
| PATL1 | RBBP4 |
| EXD2 | LSM14A |
| DIMT1 | TDP1 |
| RBM15 | METTL17 |
| CHD6 | PIP |
| WASF2 | OCIAD2 |
| PRC1 | SZRD1 |
| MGST1 | FAM162A |
| CYFIP1 | ATP5F1D |
| SMG7 | POLR2H |
| APC | AP2S1 |
| EPHB3 | ZNF746 |
| RFX7 | FOXK2 |
| PAWR | TAF3 |
| EIF3F | RNF213 |
| DHX37 | PATL1 |
| SRBD1 | EXD2 |
| TENT2 | TRIM33 |
| NUFIP1 | FUBP1 |
| NFIA | NOP2 |

| | |
|---|---|
| RARS1 | SMG7 |
| HCFC1 | SART3 |
| SNRNP27 | HDAC1 |
| CD2BP2 | CBLL1 |
| TBP | RCN2 |
| MRPL49 | PAWR |
| CNBP | WDR5 |
| TWNK | WDR77 |
| ARHGAP32 | GNB1 |
| DHX57 | MAP7D1 |
| RCN1 | TOP3A |
| DCAF7 | SRBD1 |
| HECA | ZNF639 |
| MUC6 | NFIA |
| TMOD3 | EPS15L1 |
| DIS3L2 | ACIN1 |
| CPNE3 | SECISBP2 |
| EIF5B | STRAP |
| RPS8 | THOC6 |
| LUZP1 | CD2BP2 |
| KIF2A | MRPL49 |
| PSIP1 | CNBP |
| PON1 | CPEB3 |
| RBM46 | PCNP |
| MRPS25 | DDIT3 |
| MED28 | TWNK |
| ZMAT5 | RBM27 |
| PON3 | HNRNPLL |
| MRPS22 | TIMM50 |
| WDR6 | ERI1 |
| SNW1 | MBD6 |
| RBBP8 | AFG2B |
| TRIM33 | WDR6 |
| PHF6 | APTX |
| MAP7 | RCL1 |
| SIN3A | NDUFA10 |
| RAP1B | RIOK2 |
| DNAJB12 | RPUSD4 |
| EXOSC1 | MRPL12 |
| NDUFB10 | NSDHL |
| RPP25 | EXOSC1 |
| MRPS11 | SSR3 |
| RC3H1 | CYFIP2 |
| SGPL1 | CPNE3 |

| | |
|---|---|
| NSUN2 | RBM6 |
| GTPBP10 | DGCR8 |
| PARD3 | NSUN2 |
| MEX3D | ARMCX3 |
| SART3 | DECR1 |
| KRR1 | RPL29 |
| MTPAP | ZNF326 |
| PAK1IP1 | NOP56 |
| AURKAIP1 | CHP1 |
| NOP2 | AURKAIP1 |
| PPIG | IMP3 |
| TRIM28 | RPL22L1 |
| TJP2 | APOBEC3F |
| ATP5PO | NAP1L1 |
| NAP1L1 | FAM83G |
| SCAPER | FAM83B |
| PES1 | MRPS9 |
| FUBP1 | KLHL8 |
| FAM83G | CLDN3 |
| EIF4B | ARL6IP1 |
| MOGS | ARHGAP32 |
| RALB | RMI1 |
| FOXK2 | F2 |
| NUDT16L1 | NXF1 |
| ARL6IP1 | CEP43 |
| NAT10 | DNAJC13 |
| C5 | MRM3 |
| MKRN2 | RIOX1 |
| TOR4A | SLC25A3 |
| TRMT10C | SRRM2 |
| RBM22 | AXIN1 |
| RBM27 | CMAS |
| PAN3 | RAB21 |
| CMAS | TMA16 |
| JPH1 | MRPL22 |
| LZTS3 | THOC7 |
| RAB21 | SECISBP2L |
| MRPS27 | ZBTB25 |
| MRPL22 | GTPBP4 |
| RAB6A | SLC25A13 |
| THOC7 | HERC2 |
| RBM15B | KCNQ1 |
| ZNF281 | RBM42 |
| LZTS2 | SLC39A7 |

| | |
|---|---|
| ZBTB25 | CHMP4B |
| MYBBP1A | C8orf33 |
| GFAP | EXOSC5 |
| GPATCH4 | PARD3 |
| SLX9 | THOC1 |
| CHMP4B | LUC7L3 |
| SYNGR2 | SND1 |
| EXOSC5 | KNOP1 |
| VPS28 | EBP |
| KLF16 | TMEM109 |
| RBBP6 | C1QC |
| SPTY2D1 | GSTA1 |
| ESRP1 | HCFC1 |
| THOC1 | PARP12 |
| CEMIP | SERPINC1 |
| NKRF | KIF2A |
| RNF214 | CPEB4 |
| EBP | RBM15 |
| EXOSC4 | ZNF608 |
| TAF1 | DHCR7 |
| VAPA | TRAF4 |
| C3 | EIF6 |
| PRPF19 | TMCO1 |
| EIF6 | ABI1 |
| TMCO1 | ALDH3A2 |
| TRIM56 | LTV1 |
| AFG2B | RPL14 |
| EPB41L5 | KRI1 |
| NFIX | ASCC3 |
| LTV1 | - |
| PDCD7 | TENT2 |
| POLRMT | PWP1 |
| ZNF639 | MRPL16 |
| PNO1 | MRPS7 |
| MRPL16 | CYP2S1 |
| MRPS7 | TIMM21 |
| ZCCHC17 | PFKM |
| KLK7 | BAIAP2L1 |
| NFIC | LEMD2 |
| DNAJC13 | KPNA2 |
| PNKP | CYFIP1 |
| TOE1 | FRG1 |
| YME1L1 | VRK2 |
| FLII | GATAD1 |

| | |
|---|---|
| VRK2 | RFX7 |
| MFAP4 | TNRC6A |
| TMEM70 | NOP58 |
| MTREX | NDUFS3 |
| NOP58 | RBM46 |
| SCO2 | BIN3 |
| MED4 | UBP1 |
| TOP2B | ZNF768 |
| CBX4 | SURF4 |
| HPSE | R3HDM4 |
| ZNF479 | MAP4 |
| AXIN2 | TAF1 |
| CDC5L | TOP2B |
| KIF21A | SNW1 |
| AGAP1 | CSTF2 |
| TENT4B | CEMIP |
| SMG6 | TSR1 |
| RFC1 | RBM5 |
| SPICE1 | POLR2C |
| TIMMDC1 | C3 |
| CMSS1 | KTN1 |
| REPIN1 | C5 |
| PDZD8 | AGAP1 |
| ZNF608 | RC3H1 |
| TMEM200B | SMG6 |
| CCDC137 | PAN3 |
| NOP56 | AHR |
| ZNF598 | PRPF38B |
| ZNF346 | SGPL1 |
| PPIE | NOL6 |
| SLC25A21 | TMEM45B |
| MAIP1 | SDCBP |
| | REPIN1 |
| | NMNAT1 |
| | SSR1 |
| | RBM15B |
| | MEX3D |
| | LAMB3 |
| | ZNF512 |
| | LPIN1 |
| | MRPS2 |
| | ZNF346 |
| | PPIE |
| | NAPA |

MRPL2
EPB41L4B
POLR2B
EXOSC10
