## Supplementary Table 2-7 for "Stress-Responsive Protein IFRD1 Protects Assembled Ribosomes via a Ribosome-Salvaging Mechanism"

**Supplementary Table 2. List of mouse strains used in this study**

| | | |
|---|---|---|
| B6.129-*Bhlha15*$^{tm3(cre/ERT2)Skz}$/J | Jackson laboratories | RRID: IMSR_JAX: 029228 |
| B6.Cg-*Gt(ROSA)26Sor*$^{tm9(CAG-tdTomato)Hze}$/J | Jackson laboratories | RRID: IMSR_JAX:007909 |
| Nat10 flox/flox | Generated by the authors | PMID: 40081569 |
| Ifrd1 flox/flox | RIKEN (Accession No. CDB0833K) | PMID: 26391411 |
| *Ifrd1*$^{tm1Lah}$ | Generated by Dr. Lukas Huber Medical University of Innsbruck | RRID: MGI:3043558 |
| C57BL/6J | Jackson laboratories | RRID: IMSR_JAX:000664 |

**Supplementary Table 3. List of short hairpin RNAs (shRNAs) and guide RNAs (gRNAs) for knockdown or knockout experiments.**

| Oligo | Sequence |
|---|---|
| shRNA for IFRD1_Forward | ACCGGGCAGTGGTTATAGCGATCCTTCTCGAGAAGGATCGCTATAACCACTGC |
| shRNA for IFRD1_Reverse | AAAAGCAGTGGTTATAGCGATCCTTCTCGAGAAGGATCGCTATAACCACTGCC |
| gRNA for IFRD1 KO #1_Forward | CACCGTCCTTGACGCAGATGCTTA |
| gRNA for IFRD1 KO #1_Reverse | AAACTAAGCATCTGCGTCAAGGAC |
| gRNA for IFRD1 KO #2_Forward | CACCCAATCAATATACATGCGTTC |
| gRNA for IFRD1 KO #2_Reverse | AAACGAACGCATGTATATTGATTG |

gRNA: Guide RNA

**Supplementary Table 4. List of primary antibodies used in this study**

| Name | Company | Species | Cat. No. | Purpose | Dilution |
|---|---|---|---|---|---|
| IFRD1 | Abcam | Rabbit | ab229720 | WB (Hs, Ms)<br>IP (Hs) | 1:1,000<br>5μL |
| IFRD1 | Novus | Rabbit | NBP1-87327 | WB (Ms) | 1:1,000 |
| IFRD1 | Proteintech | Rabbit | 12939-1-AP | WB (Hs) | 1:2,000 |
| GFP | Abcam | Rabbit | Ab6556 | WB | 1:1,000 |
| NAT10 | Abcam | Rabbit | ab194297 | WB | 1:1,000 |
| NCL | Cell Signaling | Rabbit | #14574 | IF | 1:200 |
| UBTF | Cell Signaling | Rabbit | #35663 | WB | 1:1,000 |
| TCOF1 | Sigma | Rabbit | HPA038237 | IF | 1:200 |
| RPS6 | Cell Signaling | Rabbit | #2217 | WB<br>IF | 1:2,000<br>1:200 |
| Phospho-S6 Ribosomal Protein (Ser240/244) | Cell Signaling | Rabbit | #5364 | WB | 1:2,500 |
| 4E-BP1 | Cell Signaling | Rabbit | #9644 | WB | 1:1,000 |
| Phospho 4E-BP1 (Thr37/46) | Cell Signaling | Rabbit | #2855 | WB | 1:1,000 |
| Acetyl-CoA Carboxylase | Cell Signaling | Rabbit | #3676 | WB | 1:1,000 |
| Phospho-Acetyl-CoA Carboxylase (Ser79) | Cell Signaling | Rabbit | #11818 | WB | 1:1,000 |
| β-Actin | Santa Cruz | Mouse | sc-47778 | WB | 1:5,000 |
| ADAR1 | Cell Signaling | Rabbit | #14175 | WB | 1:1,000 |
| dsRNA (J2) | Cell Signaling | Mouse | #76651 | IP | 5μL |
| dsRNA (K1) | Cell Signaling | Mouse | #28764 | IP<br>IF | 5μL<br>1:200 |
| ZFP36L1 | Cell Signaling | Rabbit | #30894 | WB<br>IP | 1:1,000<br>5uL |
| Ubiquitin | Santa Cruz | Mouse | sc-8017 | WB | 1:1,000 |
| G3BP1 | Abcam | Mouse | ab56574 | IF | 1:200 |
| G3BP1 | Proteintech | Rabbit | 13057-2-AP | IF | 1:200 |
| LC3B | Cell Signaling | Rabbit | #43566 | WB<br>IF | 1:1,000<br>1:200 |
| LC3B | Novus | Rabbit | NB100-2220 | WB | 1:1,000 |

| | | | | | |
|---|---|---|---|---|---|
| SQSTM1 | Santa Cruz | Mouse | sc-28359 | WB<br>IF | 1:1,000<br>1:200 |
| RPL11 | Abcam | Rabbit | ab79352 | WB<br>IF<br>IHC<br>IP | 1:2,000<br>1:200<br>1:200<br>2μg |
| RPL13 | Abcam | Rabbit | ab134961 | WB<br>IF | 1:1,000<br>1:200 |
| HA-tag | Cell Signaling | Rabbit | #3724 | WB | 1:1,000 |
| Purified anti-HA.11 Epitope Tag | BioLegend | Mouse | #901501 | IF | 1:500 |
| RPL5 | Abcam | Rabbit | ab86863 | WB<br>IF | 1:2000<br>1:200 |
| eIF2alpha | Cell Signaling | Rabbit | #5324 | WB<br>IP | 1:1,000<br>5μL |
| Phospho-eIF2alpha (Ser 35) | Cell Signaling | Rabbit | #3597 | WB | 1:1,000 |
| RPL26 | Abcam | Goat | ab157111 | IF | 1:200 |
| RPL26 | Cell Signaling | Rabbit | #2065 | WB | 1:1,000 |
| SRP68 | Abcam | Rabbit | ab157120 | WB<br>IP | 1:1000<br>5uL |
| PERK | Cell Signaling | Rabbit | #3192 | WB | 1:1000 |
| SRPR | Proteintech | Rabbit | 12090-1-AP | WB | 1:1000 |
| GAPDH | Cell Signaling | Rabbit | #2118 | WB | 1:1000 |
| α/β-tubulin | Cell Signaling | Rabbit | #2148 | WB | 1:2,000 |
| TRAPα | Santa Cruz | Mouse | sc-373916 | IF | 1:200 |
| LAMP1 | Cell Signaling | Rabbit | #99437 | IF | 1:200 |
| Amylase | Cell Signaling | Rabbit | # 3796 | IHC<br>IF | 1:500<br>1:500 |
| Puromycin | Millipore | Mouse | clone 12D10, MABE343 | WB | 1:2,000 |
| TdTomato | SicGen | Goat | AB8181 | IF | 1:200 |
| Ki-67 | Invitrogen | Rat | 14-5698-82 | IF | 1:500 |
| p-Histone H3 | Cell Signaling | Rabbit | #9701 | IHC | 1:500 |

| | | | | | |
|---|---|---|---|---|---|
| Insulin | Dako | Guinea Pig | A056401-2 | IF | 1:500 |
| E-cadherin | Cell Signaling | Mouse | #14472 | IF | 1:200 |
| CK19 | Proteintech | Rabbit | 10712-1-AP | IF | 1:200 |

WB; Western blot, IF; immunofluorescence, IHC; immunohistochemistry, IP; immunoprecipitation

**Supplementary Table 5. List of qRT-PCR primers used in this study**

| qRT-PCR primers | Species | Sequence |
|---|---|---|
| *IFRD1* Forward | Homo Sapiens | TGCAGTGGTTATAGCGATCCT |
| *IFRD1* Reverse | Homo Sapiens | CCTTGTCTTCGCACTCTTATCC |
| *Ifrd1* Forward | Mus Musculus | GTCGCATCTGTTCTTTGTATTCAG |
| *Ifrd1* Reverse | Mus Musculus | ACAGCAAACACCAAAGCAAG |
| *GAPDH* Forward | Homo Sapiens | AAGAAGGTGGTGAAGCAGGC |
| *GAPDH* Reverse | Homo Sapiens | TCCACCACCCTGTTGCTGTA |
| *Gapdh* Forward | Mus Musculus | GGGTGTGAACCACGAGAAATA |
| *Gapdh* Reverse | Mus Musculus | AGTGATGGCATGGACTGTG |
| *RPL5* Forward | Homo Sapiens | GGTGTGAAGGTTGGCCTGAC |
| *RPL5* Reverse | Homo Sapiens | GGCACCTGGCTGACCATCAA |
| *RPL11* Forward | Homo Sapiens | TCCACTGCACAGTTCGAGGG |
| *RPL11* Reverse | Homo Sapiens | AAACCTGGCCTACCCAGCAC |
| *RPS6* Forward | Homo Sapiens | TGGACGATGAACGCAAACTTC |
| *RPS6* Reverse | Homo Sapiens | TTCGGACCACATAACCCTTCC |
| 45s pre-rRNA Forward | Homo Sapiens | ACCCACCCTCGGTGAGA |
| 45s pre-rRNA Reverse | Homo Sapiens | CAAGGCACGCCTCTCAGAT |
| *SQSTM1* Forward #1 | Homo Sapiens | GCACCCCAATGTGATCTGC |
| *SQSTM1* Reverse #1 | Homo Sapiens | CGCTACACAAGTCGTAGTCTGG |
| *SQSTM1* Forward #2 | Homo Sapiens | GACTACGACTTGTGTAGCGTC |
| *SQSTM1* Reverse #2 | Homo Sapiens | AGTGTCCGTGTTTCACCTTCC |
| *TFEB* Forward | Homo Sapiens | ACCTGTCCGAGACCTATGGG |
| *TFEB* Reverse | Homo Sapiens | CGTCCAGACGCATAATGTTGTC |

**Supplementary Table 6. List of plasmid constructs used in this study**

| | | |
|---|---|---|
| IFRD1 | IFRD1 (NM_001007245) Human Tagged Lenti ORF Clone | OriGENE |
| (N)Td-Tomato-IFRD1 (Hyperactive Piggybac Transposase) | Generated by Dr. Jeffrey W. Brown | |
| (C)IFRD1- HA- GFP (Hyperactive Piggybac Transposase) | Generated by Dr. Jeffrey W. Brown | |
| IFRD1- HA (Hyperactive Piggybac Transposase) | Generated by Dr. Jeffrey W. Brown | |
| hIFRD1-His (Hyperactive Piggybac Transposase) | Generated by Dr. Jeffrey W. Brown | |
| Transposase (Hyperactive Piggybac Transposase) | Generated by Dr. Jeffrey W. Brown | |
| ΔIFRD1 | EIGC, Emory University, Dr. Laur Oskar | |

**Supplementary Table 7. List of reagents used in this study**

| | | |
|---|---|---|
| Agar | Lamda Biotech | Cat# C110 |
| Tris Buffered Saline, with Tween® 20, pH 8.0 | Sigma-Aldrich | Cat# T9039 |
| Triton X-100 | LabChem | Cat# LC262801 |
| PBS, 10X Sterile | Corning | Cat# 46-013-CM |
| Digitonin, High Purity | Sigma-Aldrich | 300410-250MG |
| Tris Buffered Saline, with Tween® 20, pH 8.0 | Sigma-Aldrich | Cat# P9039 |
| Triton X-100 | LabChem | Cat# LC262801 |
| Tamoxifen | Toronto Research Chemicals Inc | Cat# T00600 |
| Puromycin | Sigma | Cat# P8833 |
| Bortezomib | Selleck Chemicals | Cat# S1013 |
| Cycloheximide | Sigma | Cat# C7698 |
| Harringtonine | Selleck Chemicals | Cat# S9063 |
| Sodium (meta) arsenite,≥90% | Sigma-Aldrich | Cat# S7400-100G |
| Tunicamycin | Sigma-Aldrich | Cat# T7765-10MG |
| Thapsigargin | Selleck | Cat# S7895 |
| Cerulein Ammonium | Bachem | Cat# 50-259-725 |
| MG-132 | Sigma-Aldrich | Cat# 474790 |
| Rabbit Reticulocyte Lysate, Nuclease-Treated | Promega | Cat# L4960 |
| Halt™ Protease Inhibitor Cocktail, EDTA-free (100X) | ThermoFisher | Cat # 78437 |
| Halt™ Protease and Phosphatase Inhibitor Cocktail, EDTA-free (100X) | ThermoFisher | Cat# 78443 |
| RIPA Lysis and Extraction Buffer | ThermoFisher | Cat# 89900 |
| Pierce™ BCA Protein Assay Kit | ThermoFisher | Cat# 23225 |
| 2-Mercaptoethanol | Sigma-Aldrich | Cat# M3148 |
| Dithiothreitol | Research Product International | D11000-10.0 |
| NuPAGE™ 4 to 12%, Bis-Tris, 1.5 mm, Mini Protein Gel | ThermoFisher | Cat# NP0335 |
| NuPAGE™ LDS Sample Buffer (4X) | ThermoFisher | Cat# NP0007 |
| Ponceau S | Sigma-Aldrich | Cat# P7170-1L |

| | | |
|---|---|---|
| BSA | Sigma-Aldrich | Cat# A7906 |
| Dynabeads Protein A | ThermoFisher | Cat# 10001D |
| ChromoTek GFP-Trap® Magnetic Particles M-270 | Proteintech | Cat# gtd |
| Peroxidase AffiniPure Donkey Anti-Rabbit IgG (H+L) | Jackson ImmunoResearch | Cat# 711-035-152 |
| Peroxidase AffiniPure Donkey Anti-Mouse IgG (H+L) | Jackson ImmunoResearch | Cat# 715-035-150 |
| SuperSignal™ West Pico PLUS Chemiluminescent Substrate | ThermoFisher | Cat# 34579 |
| Veriblot | Abcam | Cat# ab131366 |
| Pierce™ IP Lysis Buffer | ThermoFisher | Cat# 87787 |
| Normal Rabbit IgG | Cell Signaling | Cat# 2729 |
| PowerUp SYBR Green Master Mix | ThermoFisher | Cat# A25742 |
| RNase-Free DNase Set | Qiagen | Cat# 79254 |
| PrimeScript™ RT Reagent Kit | Takara Bio Inc | Cat# RR037B |
| RNeasy Mini Kit | Qiagen | Cat# 74104 |
| Direct-zol RNA Miniprep Kits | Zymo Research Corporation | Cat # R2053 |
| TRIzol™ Reagent | ThermoFisher | Cat# 15596018 |
| Paraformaldehyde 16% Aqueous Solution EM Grade | Electron Microscopy Sciences | Cat# 15710 |
| Donkey anti-Rabbit IgG (H+L) Highly Cross-Adsorbed Secondary Antibody, Alexa Fluor 594 | ThermoFisher | A-21207 |
| Donkey anti-Rabbit IgG (H+L) Highly Cross-Adsorbed Secondary Antibody, Alexa Fluor 647 | ThermoFisher | A-31573 |
| Donkey anti-Mouse IgG (H+L) Highly Cross-Adsorbed Secondary Antibody, Alexa Fluor 488 | ThermoFisher | A-21202 |
| Donkey anti-Mouse IgG (H+L) Highly Cross-Adsorbed Secondary Antibody, Alexa Fluor 594 | ThermoFisher | A-21203 |
| Donkey anti-Mouse IgG (H+L) | ThermoFisher | A-31571 |

| | | |
|---|---|---|
| Highly Cross-Adsorbed Secondary Antibody, Alexa Fluor 647 | | |
| Donkey anti-Rabbit IgG (H+L) Secondary Antibody, Alexa Fluor 488, Invitrogen | ThermoFisher | A-21206 |
| Donkey anti-Goat IgG (H+L) Secondary Antibody, Alexa Fluor 594, Invitrogen | ThermoFisher | A-11058 |
| Donkey anti Rat IgG (H+L) Highly Cross-Adsorbed Secondary Antibody, Alexa Fluor 488 | ThermoFisher | A-21208 |
| Goat anti-Guinea Pig IgG (H+L) Highly Cross-Adsorbed Secondary Antibody, Alexa Fluor™ 647 | ThermoFisher | A-21450 |
| Lectin PNA From *Arachis hypogaea* (peanut), Alexa Fluor™ 594 Conjugate | ThermoFisher | Cat# L32459 |
| Hoechst 33342 | ThermoFisher | Cat# 62249 |
| Paraformaldehyde 16% Aqueous Solution EM Grade | Electron Microscopy Sciences | Cat# 15710 |
| ProLong Gold antifade mountant with DAPI | Invitrogen | Cat# P36930 |
| Propidium Iodide | Sigma Aldrich | Cat# P4170 |
| RPMI-1640 | Gibco | Cat# 11875093 |
| DMEM, high glucose | Gibco | Cat# 11965092 |
| Fetal Bovine Serum (FBS) | Gibco | Cat# 26140079 |
| AGS | ATCC | CRL-1739 |
| LS-174T | ATCC | CL-188 |
| HEK-293T | ATCC | CRL-1573 |
| Primocin | InvivoGen | Cat# ant-pm-1,2 |
| Corning® 100 mL Penicillin-Streptomycin Solution, 100x | Corning | Cat# 30-002-CI |
| trypLE™ Express | Gibco | Cat# 12605028 |
| RNaseOUT™ Recombinant Ribonuclease Inhibitor | ThermoFisher | Cat# 10777019 |
| 14 mL, Sterile + Certified Free | Beckman Coulter | Cat# C14302 |

| | | |
|---|---|---|
| Open-Top Thinwall Polypropylene Tube, 14 x 95mm | | |
| Sucrose | Sigma | Cat# S0389 |
| Isoflurane | Covetrus | Cat# 11695067772 |
| T-PER Tissue Protein Extraction Reagent | ThermoFisher | Cat# 78510 |
| Formaldehyde solution | Sigma-Aldrich | Cat# 252549 |
| Histo-Clear | National Diagnostics | Cat# HS-200 |
| DAB Substrate Kit | ThermoFisher | Cat# 36000 |
| Vectastain Elite ABC HRP Kit | Vector Laboratories | Cat# PK-6100 |
| Permount Mounting Medium | ThermoFisher | Cat# SP15-100 |
| Opti-MEM™ I Reduced Serum Medium | Gibco | Cat# 31985070 |
| Lipofectamine™ 2000 Transfection Reagent | Invitrogen™ | Cat# 11668019 |
| ChemiDoc™ MP Imaging System | Bio-Rad | Cat# 12003154 |
| Olympus IX83 Motorized Inverted Microscope | Olympus | |
| Odyssey Fc Blot Imaging System | LI-COR | |
| Pannoramic MIDI II | Epredia | |
| DM6B Upright Fluorescent Microscope | Leica | |
| BioTek Synergy 2 Microplate Reader | BioTek | |
| AX R Confocal System with Eclipse Ti2-E Inverted Microscope | Nikon | |
| ImageJ | NIH | https://imagej.net/ |
| Adobe Illustrator 2025 | Adobe | |
| Photoshop 2025 | Adobe | |
| SW 41 Ti Swining-Bucket Rotor | Beckman Coulter | Cat# 331362 |
| Beckman Optima LE-80K Ultracentrifuge | Beckman Coulter | Cat# 365668 |
| Piston Gradient Fractionator | BioComp | |
| GILSON Fraction Collection | Gilson | FC-203B |

| | | |
|---|---|---|
| System | | |
| Prism 10 | GraphPad | https://www. graphpad.com/scientific-software/prism/ |
| PANTHER | | https://pantherdb.org/ |
