## Supplementary material for "Stress-Responsive Protein IFRD1 Protects Assembled Ribosomes via a Ribosome-Salvaging Mechanism": Figure Legends

### Figure 1. IFRD1 is a general injury-responsive protein in vivo and in vitro

(A–C) A reduction in puromycin incorporation (A) is observed after cerulein treatment in acinar cells. (B) Without injury, the puromycin signal is stronger in acinar cells (PNA-positive) compared to ductal cells (CK19-positive) or islet cells (INS-positive) (Scale bars: top, 20 μm; bottom, 25 μm). (C) A reduction in puromycin signal in acinar cells is noted following injury (Scale bar: 20 μm). PNA: Peanut Agglutinin, INS: Insulin. (D–F) An increase in the p-eIF2α/eIF2α ratio is observed within 30 minutes of cerulein treatment in the pancreas (D), whereas IFRD1 levels remain unchanged. Red arrow indicates the IFRD1 band. Asterisk indicates a nonspecific band. At the 1-hour time point, IFRD1 levels increase at both the protein (E, left and middle) and mRNA (E, right) levels, while the p-eIF2α/eIF2α ratio returns to baseline (F). (G) X-gal staining shows IFRD1 expression (blue) exclusively in acinar cells (red box), while islets (green) and ducts (yellow) are spared. Fast Red staining was used to stain nuclei (G; Scale bars: 100 μm; inset: 20 μm). (H) Graphical overview of the cerulein injection scheme and the three stages of paligenosis. (I) Western blot of IFRD1 during the cerulein time course, with α/β-tubulin as a loading control. (J) Comparison of control and *Ifrd1*$^{\Delta/\Delta}$ mice treated with cerulein for 5 days. Cells that underwent recombination were positive for the TdTomato signal during lineage tracing and were quantified for proliferation indices: phosphorylated Histone H3 (pHH3) for cells in M phase and Ki-67 for the entire population of cycling cells. For the immunofluorescence images, the left panels use CDH1 (E-cadherin) to delineate cell boundaries, while the right panels use AMY1 (Amylase) to mark acinar cells. Images were captured at 20X magnification (Scale bar: 25 μm). (K) Time-course analysis of IFRD1 protein levels in mice conditionally knocked out for the *Nat10* gene over 28 days. (L–M) qRT-PCR (L) and Western blot (M) analyses of IFRD1 following tunicamycin treatment in HEK-293T cells. (N–O) qRT-PCR (N) and Western blot (O) analyses of IFRD1 following tunicamycin treatment in AGS cells. Asterisk indicates a nonspecific band. (P) Western blot analysis following extended tunicamycin treatment in AGS cells up to 21 hours. (Q) Western blot analysis of IFRD1 up to 4-hour tunicamycin treatment in LS-174T cells. Asterisk indicates a nonspecific band. (R) Time-course analysis of IFRD1 protein levels in AGS cells treated with 10 μM of MG-132 for up to 5 hours. Asterisk indicates a nonspecific band. For all Western blots, β-actin served as the loading control unless otherwise specified. Data are presented as mean ± standard deviation (SD) from at least three independent biological replicates. Statistical significance was defined as follows: ns, $P > 0.05$; *, $P \leq 0.05$; **, $P \leq 0.01$; ***, $P \leq 0.001$.

**Fig. 2. IFRD1 is an 80S monosome-binding protein**. (A–B) PANTHER analysis of IFRD1-interacting partners identified via mass spectrometry. Samples include pulldowns using anti-IFRD1 antibody from LS174T cells treated with tunicamycin for 3.5 hours (A, left) or 20 hours (A, middle), and pulldowns using anti-GFP antibody from AGS cells stably overexpressing HA-GFP-IFRD1 (A, right). (B) Venn diagram showing the overlap of identified proteins across the three conditions (left) and PANTHER analysis of the 199 common targets (right). (C) Immunoprecipitation of endogenous IFRD1 in LS174T and AGS cells (left and middle) demonstrates its interaction with the large ribosomal subunit proteins RPL5 and RPL11, as well as the small ribosomal subunit protein RPS6. Validation of the anti-IFRD1 antibody (ab229720) shows specific recognition of overexpressed IFRD1 while showing no cross-reactivity with its paralog, IFRD2 (right). IP: Immunoprecipitation; NC: Negative control; WCL: Whole cell lysate; OE: Overexpression. (D) Co-immunoprecipitation assays in AGS cells. HA-GFP-IFRD1 was pulled down using anti-GFP and blotted for ribosomal proteins RPL11 and RPS6 (left). Reciprocal pulldowns using anti-RPL11 and anti-RPS6 antibodies confirm the ribosomal protein interaction with tagged IFRD1 (middle and right). Red arrow indicates the IFRD1 band. Red arrows indicate the IFRD1 bands. Asterisk indicates a nonspecific band. (E–F) Ribosome fractionation via sucrose gradient centrifugation in AGS cells. Endogenous IFRD1 is observed only in the monosome fraction under both tunicamycin-treated (E) and untreated (F) conditions. RPS6 and RPL11 were used to indicate the presence of small and large subunits, respectively. (G–H) Distribution of tagged IFRD1 mainly in the monosome fraction among the ribosomal fractions in AGS cells stably overexpressing TdTomato-IFRD1 (G) or HA-GFP-IFRD1 (H). Actin is shown as a non-ribosomal control in (G). OE: overexpression. (I) Fractionation profile of AGS cells transiently transfected with ΔIFRD1 (lacking the N-terminal region). IFRD1 and HA-tag signals were detected across the gradient. The monosome is marked with a red box. PC: positive control. FL: Full length.

**Figure 3. IFRD1 localizes to cytosol and is excluded from stress granules.** (A–B) IFRD1 localizes to the cytoplasm. (A) AGS cells stably overexpressing TdTomato-tagged IFRD1 show a signal restricted to the cytoplasm. (B) The TdTomato signal is also absent from the NCL-positive (nucleolin) nucleolus. (C) Subcellular fractionation confirms enrichment of IFRD1 in the cytoplasm. This pattern is distinct from histone H3 (nucleus) and ADAR1 (nucleus; p110 isoform mainly in nucleolus). (D) Immunofluorescence shows strong cytoplasmic overlap between the TdTomato-IFRD1 signal and ribosomal proteins RPS6 (top left), RPL13 (top right), RPL5 (bottom left), and RPL11 (bottom right). Scale bars: 20 μm. (E–F) IFRD1 is not enriched in stress granules. Sodium arsenite ($NaAsO_2$) treatment induces G3BP1-positive (E) or RPS6-positive (F) stress granules. IFRD1 remains diffuse in the cytoplasm and does not concentrate within these granules. Scale bars: 20 μm in E, 10 μm in F. (G–H) HA-tagged IFRD1 displays consistent cytoplasmic localization in AGS cells treated with tunicamycin (G) or thapsigargin (H). Scale bars: 10 μm. (I) ER fractionation following a tunicamycin time course (2, 4, and 6 hours) shows IFRD1 predominantly in the non-ER cytoplasmic fraction. Markers used include GAPDH (cytosol), PERK (ER), and SRPR/SRP68 (ER). (J) Co-immunofluorescence in non-injured AGS cells shows partial spatial overlap of TdTomato-IFRD1 with the ER marker TRAPα. Scale bar: 10 μm.

**Fig. 4. IFRD1 mildly attenuates global translation by binding to a pool of idling ribosomes.**

(A–B) Puromycylation assays in AGS cells. (A) Reduced laddering is observed in cycloheximide (CHX) and tunicamycin-treated samples compared to non-treated controls. (B) Puromycylation levels are not significantly altered by either IFRD1 knockout (KO, left) or HA-IFRD1 overexpression (right). Successful knockout and overexpression are confirmed by western blotting. Ponceau S served as a loading control. (C) In vitro translation assay using rabbit reticulocyte lysate and luciferase mRNA. Human recombinant IFRD1 induces a dose-dependent reduction in luminescence compared to the BSA, used as a negative control. (D–E) Ribosomal fractionation following run-off (D) or harringtonine (E) treatment. IFRD1 remains associated with the monosome peak despite the depletion of actively translating polysomes or translation initiation block. (F–G) IFRD1 does not associate with double-stranded RNA (dsRNA). (F) Immunoprecipitation using dsRNA-binding antibodies J2 or K1 pulls down known targets ZFP36L1 and ADAR1 but fails to pull down IFRD1. p110: ADAR1 p110 isoform; p150: ADAR1 p150 isoform. (G) Co-immunofluorescence in AGS cells overexpressing TdTomato-IFRD1 shows no significant co-localization with the K1 antibody signal. Scale bar: 10 μm. (H) Lack of interaction between IFRD1 and the Signal Recognition Particle (SRP). IFRD1 does not pull down SRP68 (left), and reciprocal pull-downs using anti-SRP68 or anti-ZFP36L1 antibodies do not capture IFRD1 (middle and right). IP: Immunoprecipitation; NC: Negative control; WCL: Whole cell lysate. For all quantitative data, results represent mean ± SD of at least three independent biological replicates. Statistical significance was defined as follows: ns, $P > 0.05$; *, $P \leq 0.05$; **, $P \leq 0.01$; ***, $P \leq 0.001$; ****, $P \leq 0.0001$.

**Figure 5. IFRD1 is a ribosome-salvaging protein, the absence of which leads to degradation of 80S and 60S ribosomes and RPL11 stability.**

(A) AGS cells (Control vs. IFRD1 KO (left and right) or knockdown (KD, middle)) were treated with tunicamycin for 16 hours (left and middle) or 22 hours (right) and were subjected to ribosomal fractionation via sucrose gradient ultracentrifugation. (B) Western blot (left) and quantification (right) of ribosomal proteins RPL11, RPL5, RPL13, and RPS6 in control and IFRD1 KO cells following 6 hours of tunicamycin treatment. β-actin served as a loading control. (C) qRT-PCR analysis of mRNA expression levels for *Rpl5*, *Rpl11*, and *Rps6* in control vs. IFRD1 KO cells. Expression is relative to *Gapdh*. (D) Western blot (left) and quantification (right) of ribosomal proteins in control and IFRD1 KO cells following 16 hours of tunicamycin treatment. (E) Western blot of IFRD1

KO AGS cells treated with tunicamycin for 6 hours, along with bortezomib up to 3 hours. Ponceau S served as a loading control. (F) Western blot (left) and quantification (right) of endogenous (Endo-) and TdTomato-tagged (Tdt-) IFRD1, RPL11, and RPL5 levels in control vs. IFRD1-overexpressing (IFRD1 OE) cells after 6 hours of tunicamycin treatment. (G–H) Analysis of Ribosome biogenesis-related protein levels. Western blots and quantification for UBTF, NCL, and NAT10 in control vs. IFRD1 KO cells following 6 hours (G) or 16 hours (H) of tunicamycin treatment. (I) qRT-PCR analysis of 45S pre-rRNA levels (left) in control vs. IFRD1 KO cells. (J) Western blot and quantification show UBTF and NAT10 levels in control vs. IFRD1 OE cells after 6 hours of tunicamycin treatment. For all quantitative data, results represent mean ± SD of at least three independent biological replicates. Statistical significance was defined as follows: ns, $P > 0.05$; *, $P \leq 0.05$; **, $P \leq 0.01$; ***, $P \leq 0.001$.

**Figure 6. Loss of IFRD1 impairs autophagy and mTOR signaling during stress**

(A) Western blot analysis (left) and quantification (right) of the autophagy markers SQSTM1 (p62) and LC3B-I and -II in control cells following 6 hours of tunicamycin treatment. β-actin served as the loading control. (B–C) Comparison of autophagy markers in control and IFRD1 KO cells. Western blots and quantification show levels of SQSTM1 and LC3B-II after 6 hours (B) or 16 hours (C) of tunicamycin treatment. β-actin served as a loading control. (D) Autophagic flux analysis. Control and IFRD1 KO cells were treated with tunicamycin for 6 hours in the presence (+) or absence (-) of the lysosomal inhibitor hydroxychloroquine (HCQ). Quantification of the HCQ-induced LC3B-II accumulation (ΔLC3B-II; right) shows no significant difference (ns) between control and IFRD1 KO cells. This result indicates that the rate of autophagosomal degradation (autophagic flux) remains intact during stress. (E) qRT-PCR analysis of autophagy-related genes. mRNA levels of Sqstm1 and the transcription factor Tfeb are compared between control and IFRD1 KO cells. Expression is relative to Gapdh. (F–H) Analysis of mTORC1 activity. Western blots and quantification of phosphorylated RPS6 (p-RPS6) and phosphorylated 4E-BP1 (p-4E-BP1) relative to total protein levels in Control cells treated with tunicamycin or DMSO for 6 hours (F), or in control vs. IFRD1 KO cells after 6 hours (G) or 16 hours (H) of tunicamycin treatment. p-RPS6 levels in panels G and H were normalized to total RPS6 levels in Fig. 5B and 5D, respectively. (I) Western blot and quantification of phosphorylated acetyl-CoA carboxylase (p-ACC) relative to total ACC in Control and IFRD1 KO cells after 6 hours of tunicamycin treatment. (J) Cell viability was assessed using propidium iodide (PI) fluorescence by flow cytometry. Cells treated with 0.1% Triton X-100 were used as a positive control, whereas unstained cells resuspended in 1× PBS containing 1% BSA and 1 mM EDTA were used as a negative control. (K–L) Changes in ribosomal protein levels during injury in vivo. (K) Western blot analysis of RPL26 during a cerulein injury time course in the pancreas (0 to 18 days). α/β-tubulin served as a loading control. (L)

Immunohistochemistry for RPL11 in untreated pancreas and at days 1, 3, and 5 post-cerulein treatment. Scale bars: 50 μm; inset, 20 μm). (M–N) Immunofluorescence analysis of ribosomal protein localization. (M) Co-localization of ribosomal protein RPL11 with lysosomal marker LAMP1 in chief cells of the stomach corpus 12 hours after high-dose tamoxifen treatment. Scale bar: 50 μm. (N) Co-localization of ribosomal protein RPL26 with lysosomal makers LC3B and SQSTM1 in non-treated and cerulein-treated (Day 1) pancreatic acinar cells. Scale bar: 20 μm. (O–P) Transmission electron microscopy (TEM) analysis of autophagy. (O) Representative TEM images of pancreatic acinar cells from wild-type and *Ifrd1* KO mice at Day 1 post-cerulein treatment. Autophagosomes and autolysosomes are outlined by yellow dotted lines. Scale Bar: 2μm. (P) Quantification of total autophagosome area per image shows a significant increase in autophagosomal accumulation in chief cells in the stomach corpus in *Ifrd1* KO mice compared to wild-type controls. Scale Bar: 2μm. For all quantitative data, results represent mean ± SD of at least three independent biological replicates. Statistical significance was defined as follows: ns, $P > 0.05$; *, $P \leq 0.05$; **, $P \leq 0.01$; ***, $P \leq 0.001$.

.
