## Supplementary figures and images for "Stress-Responsive Protein IFRD1 Protects Assembled Ribosomes via a Ribosome-Salvaging Mechanism"

### Graphical Abstract

#### Ribosome Salvaging

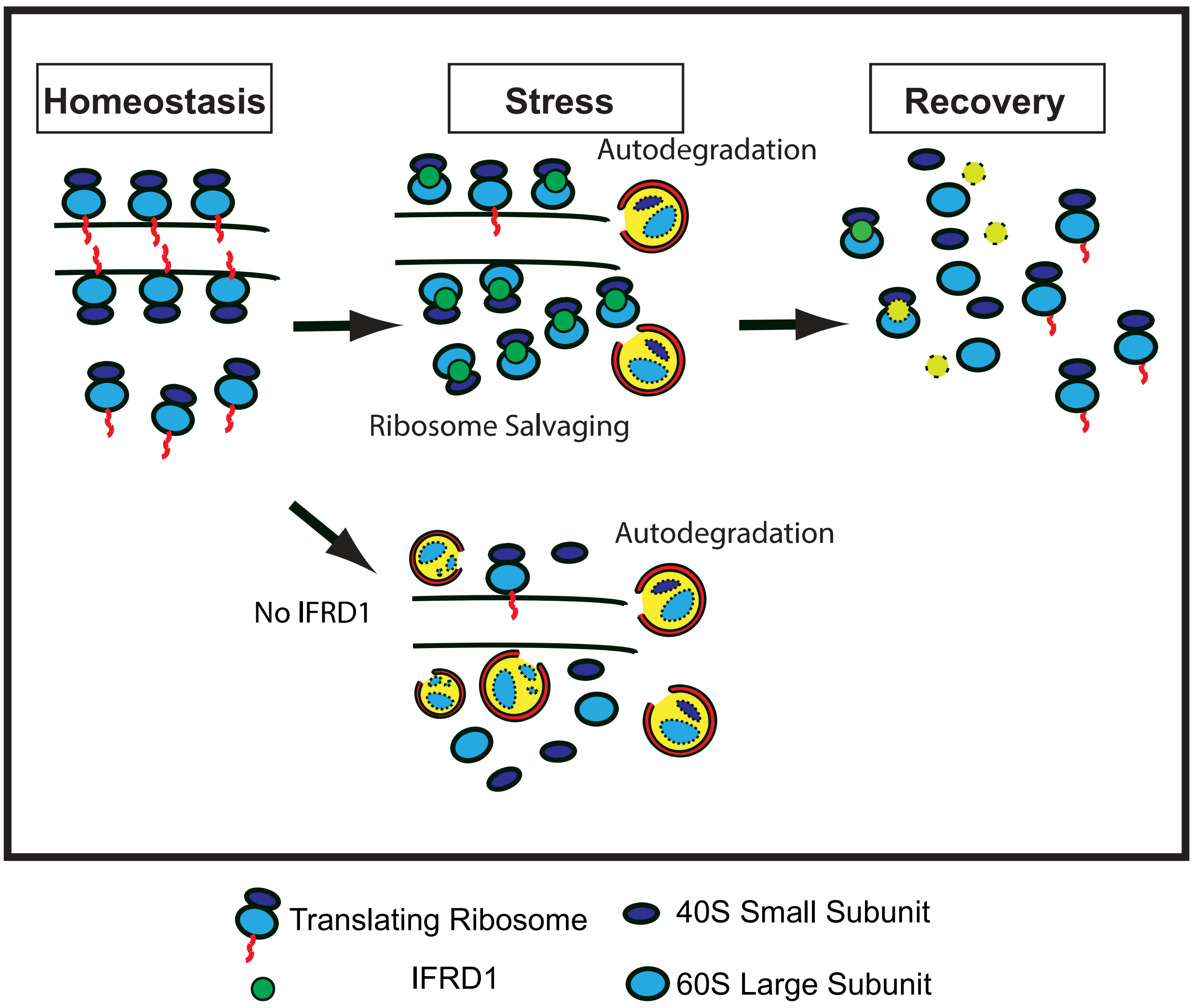
